# Domain Adaptation Improves Cross-site Classification of First-episode Schizophrenia from Multimodal Neuroimaging Data

**DOI:** 10.64898/2026.02.02.702775

**Authors:** Dominik Klepl, Barbora Rehák Bučková, Jakub Svoboda, David Tomeček, Filip Španiel, Jaroslav Hlinka

## Abstract

Identifying robust neuroimaging markers associated with schizophrenia is essential for advancing research and informing clinical understanding. However, a major obstacle to clinical translation is the limited ability of neuroimaging-based classification models to generalize across scanning sites. In this study, we first establish best performing within-site models, and then systematically investigate cross-site generalization in first-episode schizophrenia (FES) classification and evaluate strategies for mitigating site-related distribution shifts. Using data from two acquisition sites (*n* = 544), we examine ComBat, CovBat, optimal transport, and deep adaptation. Across functional, structural, and diffusion-based features, both traditional machine learning (TML) and neural network (NN) models achieved comparable performance in within-site classification, with resting state fMRI functional connectivity providing the most robust unimodal features. When models are transferred across sites, performance degraded substantially across all approaches, by 5 to 12 percentage points of balanced accuracy. Distribution-alignment methods partially mitigated this degradation. Harmonization recovered 6 to 8 points for functional connectivity in one transfer direction, mainly by correcting a displaced decision threshold. Deep adaptation recovered similar gains but less consistently. Increasing model complexity alone did not result in systematic performance gains. Across the classifiers tested on this pair of sites, multimodal feature fusion did not consistently outperform functional connectivity alone. Overall, our findings indicate that controlling for site effects was more critical than model complexity for achieving generalizable classification in FES. These results come from two sites of a single coordinated study, and whether they hold for more heterogeneous sites remains to be tested.

## 1 Introduction

Schizophrenia is a severe mental disorder characterized by marked clinical and neurobiological heterogeneity, encompassing positive symptoms such as psychosis and delusions and negative symptoms including blunted affect and social withdrawal Schultz et al. [2007]. Neuroimaging studies have variably described this heterogeneity in terms of discrete subtypes Dwyer et al. [2023] or continuous latent dimensions Kirschner et al. [2020], McWhinney et al. [2024]. Improving early detection and characterization, particularly at first episode, is therefore critical. However, current diagnostic practice relies on clinician-conducted interviews that are resource-intensive and, to a degree, subjective, motivating the development of objective neuroimaging-based classification approaches to support clinical research and understanding.

Despite the clinical importance of early-stage diagnosis, most neuroimaging-based classification studies in schizophrenia have focused on chronic patient populations, likely reflecting pragmatic constraints related to data availability. The relatively few studies targeting first-episode schizophrenia (FES) report limited reliability and poor generalization, particularly under cross-site transfer Zhuang et al. [2019], Vieira et al. [2020]. Methodological shortcomings, including non-nested cross-validation and overlap between model selection and evaluation data, have further contributed to optimistic performance estimates and limited reproducibility Vieira et al. [2020], Winterburn et al. [2019], Porter et al. [2023]. As a result, robust cross-site generalization in FES remains an open and underexplored problem with direct implications for clinical translation.

Both traditional machine learning (TML) and neural network (NN) approaches have been widely applied to structural and functional neuroimaging data, typically achieving moderate but variable performance Chu et al. [2018], Yamamoto et al. [2020b], Mahmood et al. [2019]. However, NN models are often assumed to require substantially larger sample sizes to be effective, raising questions about their practical utility in neuroimaging settings where data are inherently limited. Several studies have examined cross-site generalization in schizophrenia, predominantly in chronic or mixed-stage cohorts. These studies consistently report substantial performance degradation when models are evaluated across scanners or acquisition protocols, despite reasonable within-site accuracy Orban et al. [2018], Zeng et al. [2018], Yamamoto et al. [2020a]. Limited cross-site robustness has therefore been identified as a major barrier to clinical relevance Li et al. [2025], Porter et al. [2023]. Although domain adaptation (DA) strategies have shown promise in related neuropsychiatric conditions such as autism spectrum disorder and major depressive disorder Li et al. [2018], Fang et al. [2023], their utility for FES classification remains largely unexplored.

Evidence regarding the benefit of multimodal neuroimaging for schizophrenia classification is similarly mixed. While some studies report modest gains from combining functional, structural, diffusion, or genetic data Kanyal et al. [2024], large-scale reviews suggest little consistent advantage over unimodal approaches, with functional magnetic resonance imaging (fMRI)-derived features often emerging as the most informative Porter et al. [2023], Salvador et al. [2019]. In particular, functional connectivity (FC) has repeatedly demonstrated robust discriminative performance relative to lower-dimensional functional summaries Shevchenko et al. [2025].

In this study, we systematically compare TML and NN models for classifying individuals with FES using multimodal neuroimaging features, including fMRI, voxel-based morphometry (VBM), and diffusion-weighted imaging (DWI). Using data from two acquisition sites within the same study, we employ a rigorous nested cross-validation framework to establish within-site baselines, quantify performance degradation under cross-site transfer, and evaluate whether distribution-alignment strategies can mitigate scanner-related domain shift. This design enables a controlled assessment of model class, feature representation, and alignment strategy under conditions relevant to clinical generalization. An overview of the experimental pipeline is provided in Fig. 1.

**Figure 1:**
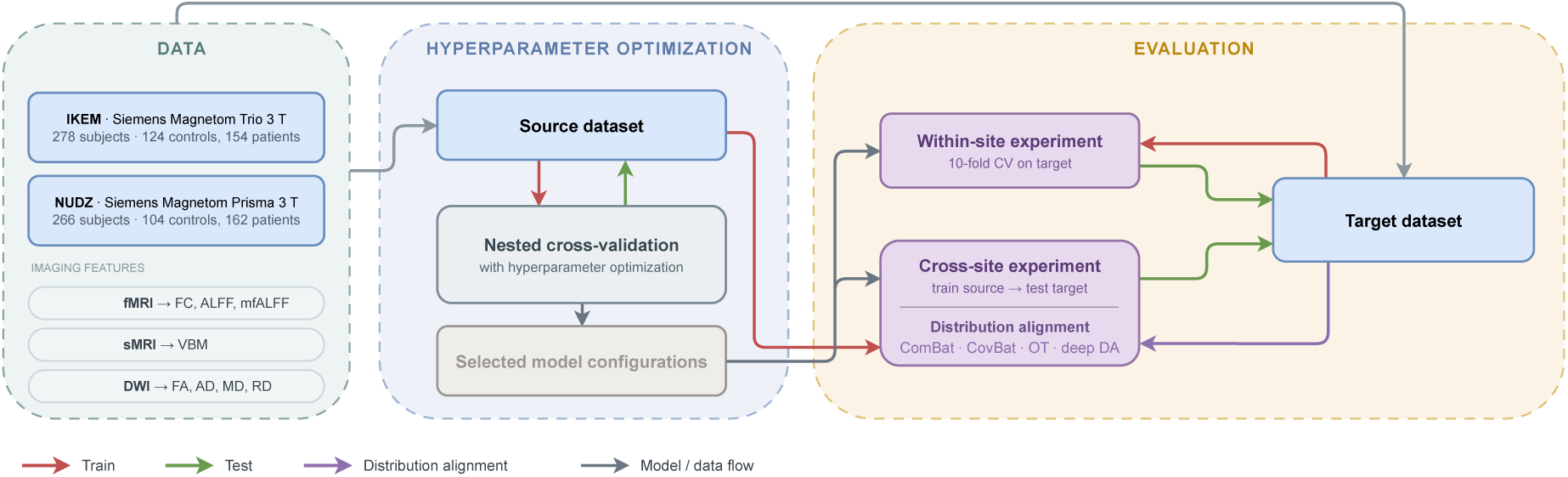
Overview of the experimental pipeline. Resting-state fMRI, T1w, and diffusion data from two acquisition sites (*N* = 544) are processed as shown. Features comprise functional connectivity (FC), regional BOLD amplitude (ALFF and its mean-normalized fractional form mfALFF), modulated gray-matter volume (VBM), and tensor metrics (FA, AD, MD, RD). Model configurations are selected by nested cross-validation on the source site. Seven classifiers are then evaluated in two paradigms. The first is within-site cross-validation on the target. The second is cross-site transfer, in which a model trained on one site is tested on the other, with and without distribution-alignment strategies.

Our primary contributions relative to the current state of the art are:

- Demonstrating that unimodal fMRI-based FC provided the most robust discrimination for FES across the two acquisition sites.
- Showing that straightforward multimodal feature fusion did not yield consistent improvements over FC alone.
- Systematically evaluating the relative performance of traditional machine-learning and neural network models under matched acquisition conditions, using a unified experimental design and rigorous nested cross-validation.
- Showing that TML approaches performed competitively with more complex NN models, while the random forest (RF) and the graph architectures ranked lowest.
- Quantifying cross-site generalization and demonstrating that feature-level harmonization and, less consistently, deep adaptation improved transfer where the site shift displaced the decision threshold, whereas optimal transport did not.

## 2 Methods

This study adopts a benchmarking framework to systematically investigate the effects of modality, model class, and site-related domain shift on classification performance and cross-site generalization. Specifically, we evaluate and compare existing methods (TML classifiers, NN architectures, and distribution-alignment strategies) within a unified experimental framework.

### 2.1 Data acquisition

Data were obtained from the Early-stage Schizophrenia Outcome (ESO) study Spaniel et al. [2016], a longitudinal clinical project conducted in the Czech Republic. Participants underwent neuroimaging at one of two acquisition sites within the same study framework: IKEM and, following a later protocol change, NUDZ. The patient group consisted of adults undergoing their first psychiatric hospitalization with a diagnosis of schizophrenia or acute and transient psychotic disorder, with untreated psychosis lasting less than 24 months; healthy control (HC) participants had no history of psychiatric disorders.

After quality control and data availability checks, the final analytic sample comprised 544 subjects (278 at IKEM, 266 at NUDZ). The exclusion criteria were applied at the level of the subject and are stated in full in Section 2.2. Demographic characteristics of each group are summarized in Table 1. All participants provided written informed consent, and the study was approved by the relevant ethics committees.

**Table 1:** Participant characteristics by site and group. Demographic characteristics of control and patient groups acquired at two sites: Institute of Clinical and Experimental Medicine (IKEM) and National Institute of Mental Health (NUDZ). Sex is reported as the number of females (F) and males (M) in each group. Age, Diagnosis to MRI, and GAF are reported as mean with standard deviation (SD) in parentheses. Diagnosis to MRI denotes the number of months elapsed between clinical diagnosis and MRI acquisition. Em dashes indicate measures not applicable to a given group. Diagnosis to MRI and GAF were recorded for a subset of the patients.

|  | N | Sex | Age | Diagnosis to MRI | GAF |
| --- | --- | --- | --- | --- | --- |
| **IKEM** |  |  |  |  |  |
| Control | 124 | 67 F, 57 M | 27.5 (SD=6.3) | — | — |
| Patient | 154 | 71 F, 83 M | 29.6 (SD=7.3) | 6.3 (SD=6.9) | 70.6 (SD=14.8) |
| **NUDZ** |  |  |  |  |  |
| Control | 104 | 63 F, 41 M | 30.1 (SD=8.0) | — | — |
| Patient | 162 | 54 F, 108 M | 26.9 (SD=6.9) | 9.1 (SD=13.9) | 64.6 (SD=15.4) |

Neuroimaging data were acquired at two sites using different scanners and acquisition protocols, reflecting hardware and protocol changes during the ESO study.

#### 2.1.1 IKEM

All data at IKEM were acquired using a Siemens Magnetom Trio 3 T scanner.

##### Structural imaging

T1-weighted MPRAGE sequence; TI = 900ms, TR = 2300ms, TE = 4.63ms, flip angle = 10°, 1 average, matrix = 256 *×* 256 *×* 224, voxel size = 1 *×* 1 *×* 1mm^3^, bandwidth = 130Hz/pixel, GRAPPA factor = 2, reference lines = 32, acquisition time = 5:30 min.

##### Functional imaging

T2-weighted echo-planar imaging (EPI) with blood oxygenation level-dependent (BOLD) contrast using SENSE imaging. TR = 2000 ms, TE = 30 ms, flip angle = 70°, 35 axial slices acquired contiguously in sequential decreasing order, voxel size = 3 × 3 × 3 mm^3^, slice dimensions = 48 × 64 voxels, 400 functional volumes. The structural image acquired at the beginning of the session was used for anatomical reference.

##### Diffusion-weighted imaging

Spin-Echo EPI sequence, TR = 8300ms, TE = 84ms, matrix = 112 × 128, voxel size = 2 × 2 × 2mm^3^, b-value = 0 and 900s/mm^2^ in 30 diffusion gradient directions, 2 averages, bandwidth = 1502Hz/pixel, GRAPPA factor = 2 (phase-encoding direction), reference lines = 24, prescan normalize = *of f*, elliptical filter = *of f*, raw filter = *on* (intensity: weak), acquisition time = 9 : 01 min.

#### 2.1.2 NUDZ

All data at NUDZ were acquired using a Siemens Magnetom Prisma 3 T scanner.

##### Structural imaging

T1-weighted MPRAGE sequence; TI = 1000ms, TR = 2400ms, TE = 2.34ms, flip angle = 8°, matrix = 320 × 320, voxel size = 0.7 × 0.7 × 0.7mm^3^, field of view = 224 × 224mm^2^.

##### Functional imaging

T2*-weighted gradient EPI sequence sensitive to BOLD contrast, TR = 2000ms, TE = 30ms, flip angle = 70°, 37 axial slices acquired contiguously in alternating increasing order, voxel size = 3 × 3 × 3mm^3^, matrix = 64 × 64, field of view = 192 × 192mm^2^, 300 functional volumes.

##### Diffusion-weighted imaging

Spin-Echo EPI sequence; TR = 8200 ms, TE = 83 ms, flip angle = 90*^◦^*, matrix = 98 × 98, 57 contiguous axial slices with slice thickness = 2 mm, voxel size = 2 × 2 × 2 mm^3^, field of view = 260 × 211.25 mm^2^, *b*-values of 0, 1100, and 2500 s/mm^2^ in 64 diffusion gradient directions.

### 2.2 Preprocessing

The functional and structural data of both acquisition sites were processed identically. Resting-state fMRI data were preprocessed with fMRIPrep 24.1.1 Esteban et al. [2019]. The workflow comprised intensity non-uniformity correction and skull stripping of the T1-weighted reference, tissue segmentation, head-motion correction of the BOLD runs, coregistration to the anatomical reference, and non-linear normalization to the MNI152NLin6Asym template Avants et al. [2008] at 2 mm isotropic resolution, which matches the resolution of the parcellations. Surface reconstruction was disabled, since all subsequent analyses are volumetric. Framewise displacement and the physiological component time series were retained from the fMRIPrep confound file for the denoising step described below.

Denoising was performed with nilearn Abraham et al. [2014] on the voxel time series. The confound model comprised 34 regressors: the six rigid-body motion parameters, their temporal derivatives, and the squares of both Friston et al. [1996], together with the five leading white-matter and the five leading cerebrospinal-fluid aCompCor components Behzadi et al. [2007]. Confound regression and temporal band-pass filtering to 0.008–0.09 Hz were carried out in a single operation. The lower cutoff was set at 0.008 Hz to match the 128 s discrete-cosine high-pass filter that fMRIPrep applies before estimating the aCompCor components.

Regional time series were obtained by averaging the denoised voxel time series within each parcel. The average was taken over voxels that were free of missing values and non-constant in time (thus excluding the zero-valued voxels outside the field of view), so that parcels only partially covered by the field of view are not biased toward zero. A parcel with no usable voxel in a given subject was recorded as missing rather than as zero.

Regional gray-matter volume was derived from the same fMRIPrep anatomical outputs, so that both modalities share one segmentation and one spatial normalization. The gray-matter probability map in template space was multiplied voxel-wise by the Jacobian determinant of the T1-weighted-to-template warp, including its affine component, and summed within each parcel. The resulting modulated volume in mm^3^, summed over the brain, was compared with each subject’s native gray-matter volume to verify the spatial transformation; their ratio was 0.96 to 1.00 (median 1.00; a value of 1 indicates a volume-preserving transformation) in all subjects.

DWI data were preprocessed using tools from FSL and MRtrix3 Tournier et al. [2019], including denoising Veraart et al. [2016], Gibbs ringing correction Kellner et al. [2016], susceptibility-induced distortion correction using images with reversed phase-encoding directions Andersson et al. [2003], Skare and Bammer [2010], and correction for motion and eddy-current distortions using eddy_cuda Andersson and Sotiropoulos [2016].

#### Quality control and exclusion criteria

Exclusions were decided per subject, on the functional run, before any modeling. A subject was retained if the run satisfied four criteria: mean framewise displacement below 0.30 mm, less than 25% of volumes exceeding 0.5 mm, a median temporal signal-to-noise ratio of at least 20, and at least 80 retained volumes. This yielded 544 subjects, 278 at IKEM and 266 at NUDZ.

### 2.3 Feature extraction

For resting-state fMRI, regional time series were extracted using a subset of the Automatic Anatomical Labeling (AAL) atlas Tzourio-Mazoyer et al. [2002] (90 cortical and subcortical regions). FC was computed as Fisher-z-transformed Pearson correlations between all pairs of regions, yielding a 90 × 90 connectivity matrix per subject. Group-averaged FC matrices for each site and diagnostic group, together with the corresponding patient-minus-control difference maps, are shown in Fig. 2. In addition, the amplitude of low-frequency fluctuations (ALFF) Yang et al. [2007] of the BOLD signal in the 0.008–0.09 Hz band was computed for each region. Because the raw amplitude is expressed in arbitrary BOLD units and carries a large scanner-dependent offset, we also computed the mean-normalized fractional amplitude of low-frequency fluctuations (mfALFF), which divides band-limited power by total spectral power and normalizes by the whole-brain mean, as the summary functional-amplitude feature throughout. To examine the sensitivity of findings to the choice of parcellation, supplementary analyses were conducted on three additional atlases (Harvard–Oxford Desikan et al. [2006], Makris et al. [2006], the Schaefer100 cortical parcellation Schaefer et al. [2018] combined with the Tian S2 subcortical atlas Tian et al. [2020], and Brainnetome246 Fan et al. [2016]) spanning 112 to 246 regions. Results are reported in the Supplementary materials.

**Figure 2:**
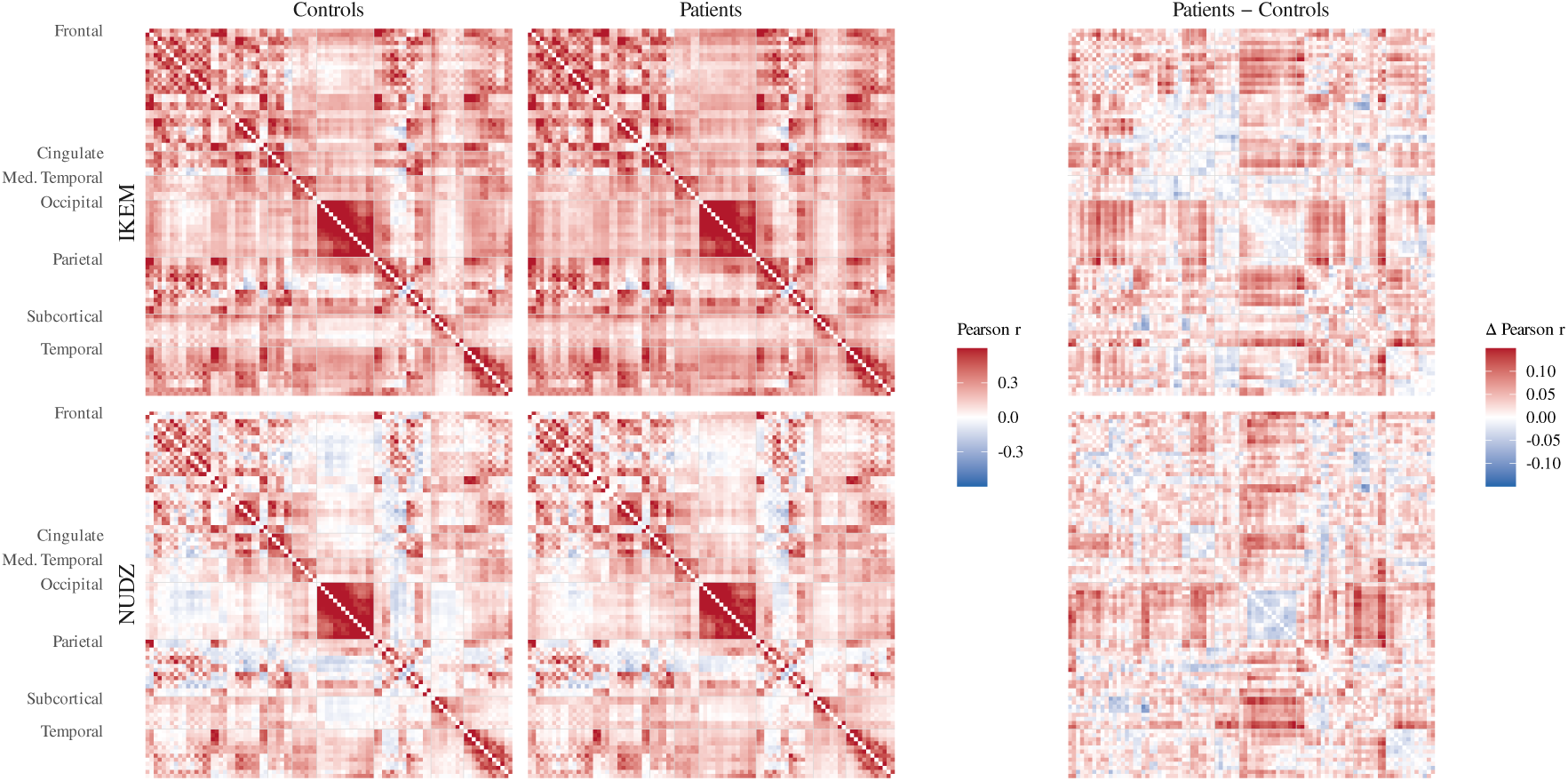
Group-average functional connectivity matrices. Pairwise connectivity averaged in Fisher-z space and back-transformed to Pearson *r*, shown separately for controls and patients at each acquisition site (IKEM, NUDZ), together with the corresponding group difference (Patients *−* Controls). Regions are ordered according to the AAL90 atlas and grouped by anatomical regions.

From structural MRI data, VBM measures were extracted at the regional level using the same parcellations, as modulated gray-matter volume in mm^3^.

For DWI, diffusion tensors were fitted voxel-wise to compute fractional anisotropy (FA) Basser et al. [1994], Westin [1997], axial diffusivity (AD), mean diffusivity (MD), and radial diffusivity (RD). All four metrics were averaged within the white-matter tracts of the JHU atlas. Because these features are defined on a tract atlas, they are independent of the gray-matter parcellation and were therefore evaluated in the AAL90 analyses only. Diffusion data were available for 344 of the 544 subjects; the diffusion-only models are evaluated on that subset, whereas in combined feature sets the missing diffusion block is imputed within each training fold.

### 2.4 Experimental design

We aimed to systematically evaluate which modeling strategies yield the most robust performance for FES classification under realistic acquisition variability. Specifically, we examined unimodal versus multimodal feature representations, TML versus NN models, and the impact of direct cross-site transfer versus distribution-alignment strategies.

Model performance was evaluated under two settings: within-site and cross-site classification. Hyperparameter optimization was performed using data from one acquisition site (the *source site*), and the resulting configurations were reused unchanged in all subsequent experiments. In the within-site setting, models were trained and evaluated using stratified 10-fold cross-validation, repeated with five different splits, on data from the other acquisition site (the *target site*), which had not been used during configuration selection, providing an unbiased estimate of performance under matched acquisition conditions. In the cross-site setting, models were trained on all available source-site data and evaluated on the target site without access to target labels, explicitly quantifying performance degradation induced by scanner- and protocol-related domain shift. Distribution-alignment strategies were evaluated exclusively in this cross-site setting. Each transfer direction is named after its target site, so that transfer to IKEM means training on NUDZ and testing on IKEM. In every setting, features were standardized and missing values imputed using statistics estimated on the training fold only.

#### 2.4.1 Hyperparameter optimization

Hyperparameters were optimized with Bayesian search using a Tree-structured Parzen Estimator under nested cross-validation with ten outer and five inner folds. The search was a single pass over the full space, with 50 trials per inner search of which the first ten were drawn at random to initialize the estimator, and the same procedure was used for the TML and the NN models. The configuration selected by the inner search in each of the ten outer folds was retained, giving ten hyperparameter configurations per source site, modality, and model class. Every subsequent analysis trained new models with these configurations rather than reusing the fitted models, and their predictions were pooled at the subject level (Section 2.8).

All selected configurations were fixed and reused unchanged in subsequent within-site and cross-site experiments. No data used for final performance evaluation were involved in hyperparameter selection.

### 2.5 Traditional machine learning models

We evaluated three widely used TML classifiers representing distinct modeling paradigms: ridge regression (RR), support vector machine (SVM), and RF. Classifier-specific hyperparameters were selected within the nested cross-validation procedure described above. For RR, the regularization strength was tuned; SVM models varied in kernel type and regularization parameters; and RF models varied in the number of trees, tree depth, and split-related parameters. Full hyperparameter ranges are reported in the Supplementary materials.

### 2.6 Neural network models

We evaluated several NN architectures previously used in neuroimaging-based classification, including a multilayer perceptron (MLP), two versions of graph neural networks (GNNs), and the row- and column-wise convolutional neural network (RowColCNN) architecture designed for connectivity matrices.

All neural models followed a standard encoder-classifier structure, where an encoder produced a latent representation that was passed to a lightweight MLP classifier. Optimization-related hyperparameters common across architectures included the learning rate, weight decay, learning-rate scheduling, and dropout. Classifier-specific hyperparameters controlled the number of layers, the hidden-layer width, and the dropout rate. When multilayer classifiers were used, hidden-layer dimensionality was progressively reduced relative to the encoder output.

Models were trained for up to 400 epochs with early stopping on a 10% validation split taken from the training data of each cross-validation fold, terminating when validation loss had not improved for 20 epochs. Full hyperparameter ranges are reported in the Supplementary materials.

#### 2.6.1 Multilayer perceptron

The MLP operated on vectorized input features and was therefore applicable to all imaging modalities. Hyperparameters included the number of layers, dropout rate, and layer dimensionalities.

#### 2.6.2 Graph neural network

We evaluated two GNN architectures: graph convolutional network (GCN) Kipf [2016] and graph attention network (GAT) Veličković et al. [2018], Brody et al. [2022]. FC matrices defined a fixed graph structure, while node-level features included regional mfALFF, rows of the connectivity matrix, and VBM-derived measures. Graph structure was fixed and not learned during training.

Hyperparameters included the number of layers, hidden dimensionality, dropout, normalization, pooling strategy, and optional jumping-knowledge mechanisms. Graph sparsification was treated as a hyperparameter using either thresholding or *k*-nearest-neighbor construction. For GAT, a binary choice between the original formulation and GATv2 was also considered.

#### 2.6.3 Row- and column-wise convolutional neural network

RowColCNN Xu et al. [2023] is a neural architecture specifically designed for symmetric matrices and was applied exclusively to FC data. The model applies successive one-dimensional convolutions along rows and columns of the connectivity matrix, followed by batch normalization, ReLU activation, and dropout. The numbers of row-wise and column-wise convolutional kernels determined the encoder output dimensionality and were treated as tunable hyperparameters.

### 2.7 Distribution-alignment and domain adaptation

To mitigate scanner- and protocol-related distribution shifts in the cross-site setting, we evaluated a range of distribution-alignment strategies that differ in their underlying mechanisms and modeling assumptions, spanning feature-level harmonization, feature-level transport, and latent-level deep adaptation. All alignment and adaptation strategies were applied exclusively in the cross-site setting and compared against unaligned baseline, and none used target-site labels. As a reference, we additionally report the within-site model, trained and evaluated on the target site by cross-validation, which approximates the achievable ceiling. All strategies were evaluated inductively. The target-site subjects were split into ten stratified folds, and for each fold the alignment was estimated from the source site and the other nine target folds only. The resulting model then classified the held-out fold, to which harmonization parameters estimated on the nine folds were applied unchanged. No target subject therefore contributed to the alignment used to classify it.

#### 2.7.1 ComBat harmonization

combining batches (ComBat) harmonization Fortin et al. [2018] models site-related effects as feature-wise additive and multiplicative biases, which are estimated within an empirical Bayes framework. The intent is to remove the site effect while preserving inter-subject variability.

In our experiments, ComBat was applied at the feature level using unlabeled data from both sites, with scanner identity treated as the batch variable and no access to diagnostic labels. Harmonized features were then used for cross-site training and evaluation. We additionally evaluated covariance batches (CovBat) Chen et al. [2022], which extends ComBat by additionally harmonizing the covariance structure across sites, under the same label-free protocol.

#### 2.7.2 Optimal transport–based alignment

As an alternative, model-agnostic strategy, we evaluated distribution alignment based on optimal transport (OT) Courty et al. [2016]. OT aligns probability distributions by estimating a transport plan that minimally shifts samples from one domain to another according to a predefined cost function, without assuming a parametric form of the domain shift. The transport plan was estimated using unlabeled data from both sites and applied to source-site features prior to classifier training, aligning them to the target-site distribution. The transport plan was computed exactly, without entropic regularization, with uniform weights on the subjects of each site, on the one-dimensional decision score of the classifier fitted on the source site. The resulting barycentric map was applied to the full feature vectors, and the classifier was refit on the transported source data with the source labels. This procedure was evaluated for all model classes and modalities in the cross-site setting.

#### 2.7.3 Deep latent-level adaptation

For NN-based models, we evaluated a range of deep domain-adaptation methods that align source- and target-site representations in latent space using unlabeled target-site data. The architectures themselves are those described in Section 2.6. This included adversarial approaches, adversarial discriminative domain adaptation (ADDA) Tzeng et al. [2017] and domain-adversarial neural network (DANN) Ganin et al. [2016], a Wasserstein-based variant, Wasserstein distance guided representation learning (WDGRL) Shen et al. [2018], and moment-matching approaches, DeepCORAL Sun and Saenko [2016] and deep adaptation network (DAN) Long et al. [2015].

The procedure consisted of two stages. First, encoder–classifier models were trained on labeled data from the source site. In the second stage, the encoder was adapted using unlabeled target-site data while keeping the classifier fixed, minimizing a method-specific measure of the discrepancy between source and target latent distributions. That discrepancy was an adversarial domain-discriminator objective for ADDA and DANN, a Wasserstein-distance critic for WDGRL, and a moment-matching penalty for DeepCORAL and DAN. After adaptation, the adapted encoder and fixed classifier were evaluated on the target site. In the main text we report ADDA as a representative deep strategy; the full panel is reported in the Supplementary materials.

### 2.8 Performance evaluation and implementation

All experiments were implemented in Python 3.11.15. TML models were implemented using scikit-learn Pedregosa et al. [2011], while NN models were implemented using PyTorch Paszke et al. [2019] and PyTorch Geometric Fey and Lenssen [2019]. Statistical analyses and visualization were performed in R 4.6.1 Team et al. [2021] using the tidyverse Wickham et al. [2019]. All experiments were run on a cluster of 16 identical nodes, each with one 32-core AMD EPYC 9374F processor and 376 GiB of memory, under Rocky Linux 9. No graphics accelerators were used; all models, including the neural architectures, were trained on CPU.

Model performance was evaluated at the subject level. For each subject, the predictions of the ten retained hyperparameter configurations of a given model class were averaged, giving one predicted class probability and one predicted label per subject, the latter obtained by majority vote over the ten predicted labels, with an exact tie broken by the averaged probability. The averaged probability was used for the area under the ROC curve. All performance metrics were computed from these pooled per-subject predictions. This approach yields a more conservative and clinically relevant assessment by approximating a realistic deployment scenario in which a single diagnostic decision is made per subject, while accounting for variability across plausible model configurations.

Balanced accuracy was used as the primary performance metric to account for class imbalance between FES and HC. Confidence intervals were estimated using non-parametric bootstrap resampling over subjects (2000 resamples). The area under the ROC curve, computed from the pooled class probabilities, together with sensitivity and specificity, is reported as a secondary measure with confidence intervals from the same bootstrap.

To compare several methods evaluated on the same subjects, each subject was scored as correct or incorrect under every method and the resulting balanced accuracies were compared with the stratified paired bootstrap described below.

Pairwise comparisons between individual classifiers were made with a stratified paired bootstrap. Subjects were resampled with replacement within each class, every method being recomputed on the same resample so that the comparison remains paired, and the two-sided *p*-value was taken as

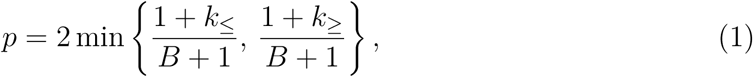

where *B* = 10 000 is the number of resamples, Δ*_b_* is the difference in balanced accuracy between the two methods in resample *b*, and *k_≤_* and *k_≥_* are the numbers of resamples with Δ*_b_ ≤* 0 and Δ*_b_ ≥* 0, respectively. Resamples in which the difference is exactly zero therefore count against significance on both sides. Besides the comparisons between all pairs of methods, each alignment strategy was tested against direct transfer and each feature combination against FC alone. Within each family of comparisons the *p*-values were adjusted by the Holm method. The same procedure was applied to the area under the ROC curve when comparing direct transfer with the within-site reference and each alignment strategy with direct transfer.

Prediction agreement between two feature representations was assessed at the level of individual subjects. For a given classifier and site, each subject was scored as correct or incorrect under each representation, and the resulting 2 × 2 table was summarized by the observed agreement and by Cohen’s *κ*. The chance level quoted alongside the observed agreement is the expected agreement of that table, computed from the marginal accuracy of the two representations under independence. It therefore varies with how accurate each representation is, rather than being fixed at 50%.

### 2.9 Decision threshold and group separation

A drop in balanced accuracy under cross-site transfer can have two causes. The site shift may move all predicted probabilities relative to the 0.5 threshold above which a subject is classified as a patient, while patients still score higher than controls. Alternatively, it may make the probabilities of the two groups overlap more, i.e. a loss of group separation. The ROC curve traces sensitivity and specificity across all thresholds, so moving the threshold moves a classifier along the curve, whereas a change in group separation moves the curve itself. For each cross-site strategy, the sensitivity and specificity at the 0.5 threshold give its operating point, and its offset is its balanced accuracy minus that of the direct-transfer ROC curve at the comparable point. The offset is zero on the curve, where only the threshold moved, positive above it, where group separation improved, and negative below it. The within-site reference was placed on the same curve, to show how much group separation direct transfer had lost. The offset and the change in the area under the ROC curve were tested with the paired bootstrap and Holm adjustment described above. Because this analysis uses the averaged probability rather than the majority vote, its balanced accuracies can differ slightly from those reported elsewhere.

## 3 Results

We first report results from within-site classification experiments to establish baseline performance under matched acquisition conditions and to assess the relative contribution of imaging modalities, feature combinations, and model classes. We then present results from cross-site classification experiments, which explicitly probe generalization under domain shift and evaluate the effectiveness of distribution-alignment and domain-adaptation strategies. Results are reported as balanced accuracy, with statistical comparisons based on the stratified paired bootstrap over per-subject correctness. Exact numerical values for every figure are reported in the Supplementary materials.

### 3.1 Unimodal performance across imaging modalities

Across both acquisition sites and model classes, FC consistently achieved the highest balanced accuracy under matched acquisition conditions (Fig. 3), reaching 78.6% at IKEM and 77.4% at NUDZ for the best of the four classifiers compared here. This pattern was observed for both TML and NN approaches. In contrast, unimodal representations derived from ALFF, VBM, and DWI showed systematically lower performance and greater variability across sites, with VBM (52.9–58.3%) and DWI (45.8–55.2%) close to chance throughout.

**Figure 3:**
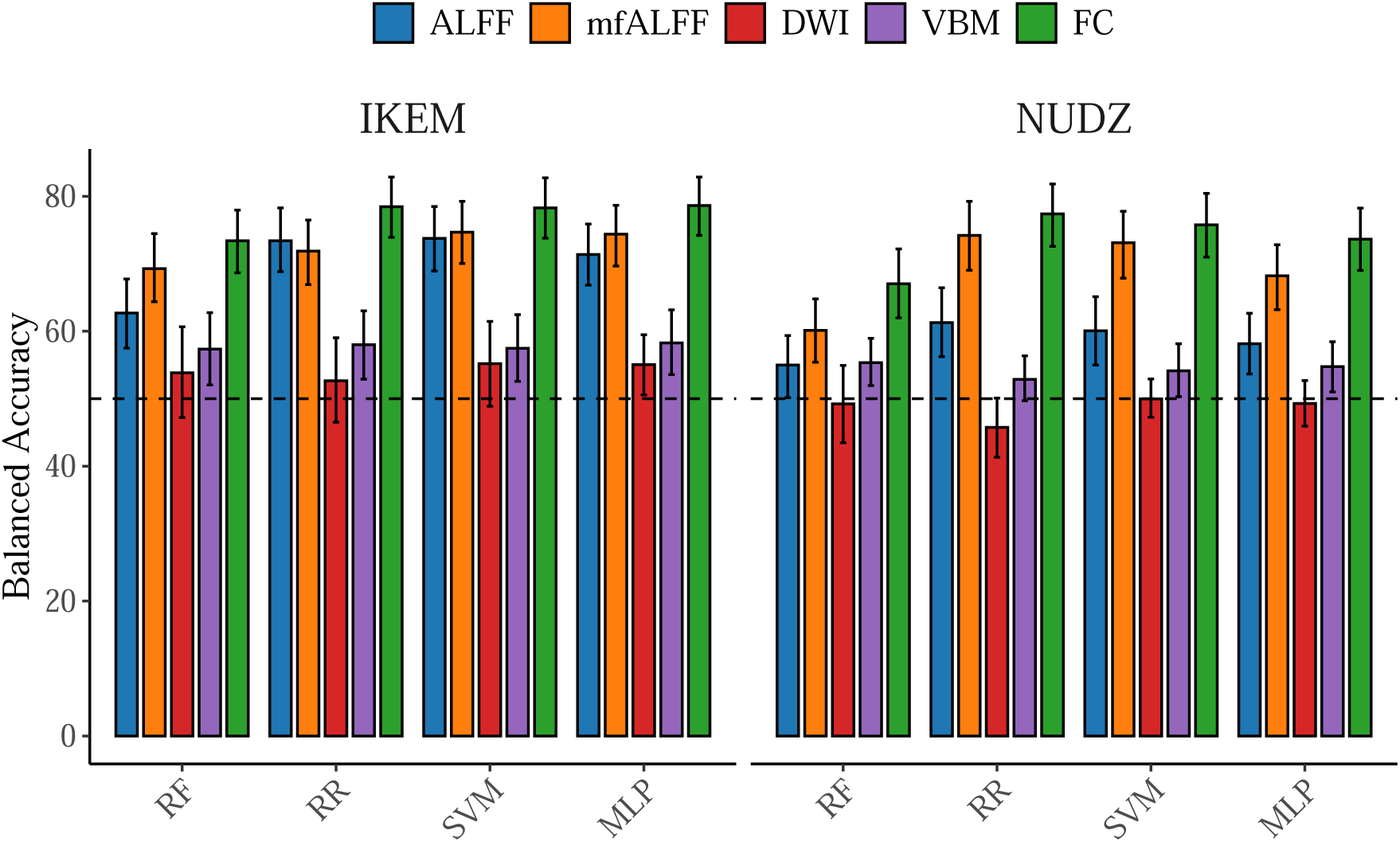
Within-site classification performance across imaging modalities. Bars show balanced accuracy (mean *±* 95% bootstrap confidence intervals; 2000 resamples) obtained in within-site cross-validation, reported separately for the IKEM and NUDZ cohorts. Results are shown for unimodal feature representations derived from resting-state fMRI amplitude of low-frequency fluctuations (ALFF) and its mean-normalized fractional variant (mfALFF), diffusion-weighted imaging (DWI), voxel-based morphometry (VBM), and functional connectivity (FC), evaluated using four representative classifiers: ridge regression (RR), support vector machines (SVM), random forests (RF), and multilayer perceptrons (MLP). The DWI bars are evaluated on the 344 subjects with diffusion data; all other bars use the full cohort. The horizontal dashed line denotes chance-level performance (50% balanced accuracy). Error bars reflect uncertainty across bootstrap resamples of subjects.

The mfALFF emerged as a strong low-dimensional alternative despite comprising only 90 regional values against the 4005 edges of the FC representation. At NUDZ, mfALFF performed close to FC (74.2% for RR against 77.4% for the same classifier on FC). The paired difference was 3.2 percentage points (*p* = 0.34) and was therefore not statistically significant. At IKEM the gap was 6.6 points for RR (*p* = 0.03) and 3.6 to 4.3 points for the other classifiers. Raw ALFF was substantially weaker than its mean-normalized form at NUDZ (61.3% against 74.2% for the best classifier of each), whereas at IKEM the two performed comparably (73.7% against 74.7%). This site-dependent discrepancy is consistent with the scanner-dependent amplitude offset that motivated the normalization.

Paired bootstrap comparisons of per-subject correctness confirmed the robustness of these descriptive patterns. The paired comparison in the Supplementary materials indicates that FC-based models achieve consistently higher balanced accuracy than alternative unimodal representations across most model–site combinations. Based on these results, FC was selected as the primary unimodal representation for subsequent analyses.

### 3.2 Effect of multimodal feature fusion

We next examined whether combining FC with additional imaging modalities improves classification performance under matched acquisition conditions (Fig. 4). Across sites and model classes, balanced accuracy was broadly similar for FC-only and multimodal feature representations.

**Figure 4:**
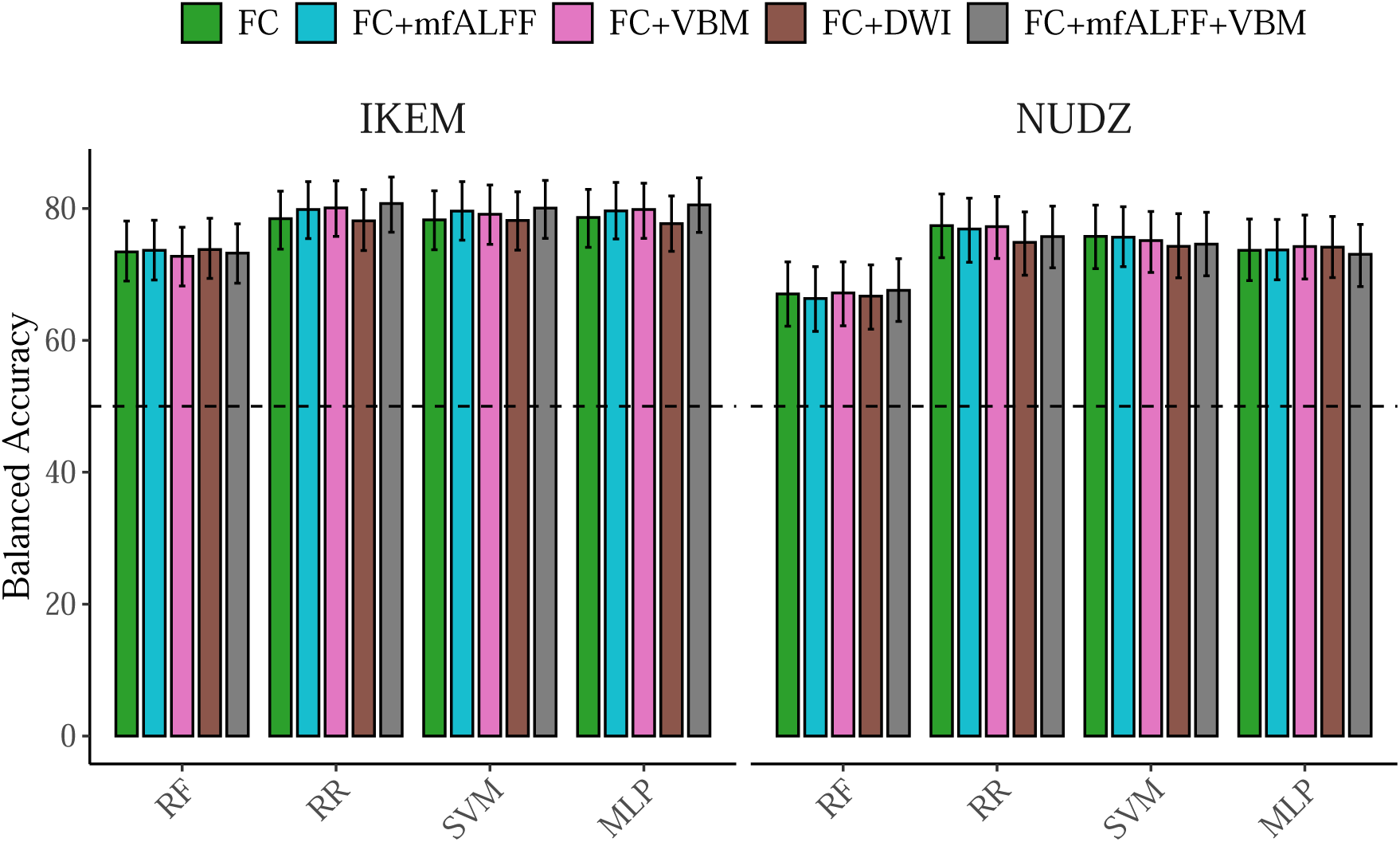
Within-site classification performance for functional connectivity and multimodal feature combinations. Bars show balanced accuracy (mean *±* 95% bootstrap confidence intervals; 2000 resamples) obtained in within-site cross-validation, reported separately for the IKEM and NUDZ cohorts. Performance is shown for functional connectivity (FC) features alone and for multimodal extensions that augment FC with additional imaging modalities, including FC+mfALFF, FC+VBM, FC+DWI, and FC+mfALFF+VBM. Results are reported across four representative classifier families: ridge regression (RR), support vector machines (SVM), random forests (RF), and multilayer perceptrons (MLP). The horizontal dashed line denotes chance-level performance (50% balanced accuracy). Error bars reflect uncertainty across bootstrap resamples of subjects.

Differences between FC and its multimodal extensions were generally small, heterogeneous, and characterized by substantial overlap of confidence intervals. Paired bootstrap comparisons against the corresponding FC-only model did not reveal any multimodal configuration that consistently outperformed FC across classifiers or sites (Supplementary materials). Across the four classifiers the deltas ranged from *−*2.5 to +2.3 percentage points. After Holm correction, no feature combination and no classifier significantly exceeded RR on FC alone. Adding diffusion features (FC+DWI) tended to reduce performance rather than improve it, significantly so for RR at NUDZ (*−*2.5 points, *p* = 0.021). FC was therefore retained as the primary representation for the subsequent model-comparison and cross-site analyses.

FC and mfALFF largely did not misclassify the same individuals. Per-subject agreement between the two classifiers, scored as correct or incorrect, ranged from 61% to 70% against chance levels of 57% to 65% (Supplementary materials) (Cohen’s *κ* 0.02 to 0.15 for seven of the eight classifier-by-site combinations, and 0.30 for the RF at NUDZ). Of the subjects misclassified from FC, 30% to 42% were also misclassified from mfALFF for seven of the eight classifier-by-site combinations, rising to 58% for the RF at NUDZ. The partially distinct classification error patterns suggest that the two representations may carry complementary predictive information.

### 3.3 Comparison of model classes under matched acquisition conditions

We compared all seven model classes under matched acquisition conditions using FC features (Fig. 5). Although aggregate balanced accuracy showed only modest absolute differences between the leading classifiers, the paired comparison revealed a consistent performance hierarchy across sites (Supplementary materials).

**Figure 5:**
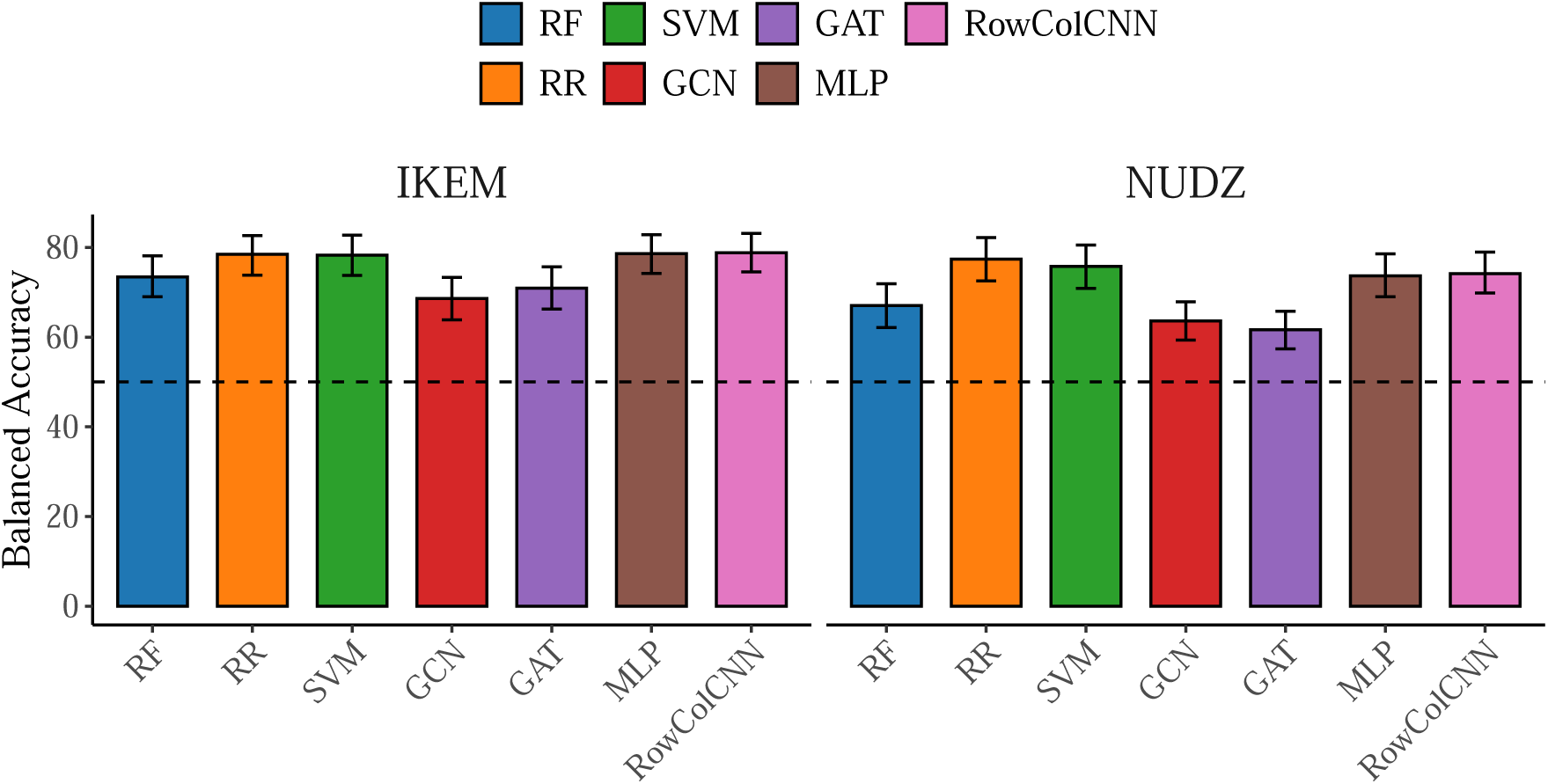
Within-site classification performance across model classes. Bars show balanced accuracy (mean *±* 95% bootstrap confidence intervals; 2000 resamples) obtained in within-site cross-validation, reported separately for the IKEM and NUDZ cohorts. Performance is shown for functional connectivity (FC) features. Results are reported for all evaluated model classes: ridge regression (RR), support vector machines (SVM), random forests (RF), multilayer perceptrons (MLP), graph convolutional networks (GCN), graph attention networks (GAT), and the Row–Column convolutional neural network (RowColCNN). The horizontal dashed line denotes chance-level performance (50% balanced accuracy). Error bars reflect uncertainty across bootstrap resamples of subjects.

Across both acquisition sites, RR achieved the highest balanced accuracy at NUDZ (77.4%) and was within 0.3 points of the best classifier at IKEM (78.5%), followed by the SVM (78.3% and 75.8%) and the MLP (78.6% and 73.6%). The RowColCNN also ranked among the strongest performers (78.8% and 74.1%). The four classifiers were mutually indistinguishable at IKEM. At NUDZ, RR exceeded the RowColCNN and the MLP by 3.3 and 3.8 points, at the margin of significance (*p* = 0.050 and *p* = 0.053). In contrast, more complex architectures such as GCN, GAT, and RF exhibited lower performance (68.6% and 63.6%, 70.9% and 61.6%, and 73.4% and 67.0%, respectively). Both GNNs fell significantly below each of the four leading classifiers at both sites, as did the RF at NUDZ. Increasing architectural complexity therefore yielded no systematic advantage.

Overall, RR, the SVM, and the MLP demonstrated stable performance at both sites, with the RowColCNN performing consistently well in the unimodal FC setting. These models were therefore selected for subsequent cross-site analyses.

### 3.4 Cross-site generalization and impact of distribution-alignment strategies

Cross-site evaluation revealed a pronounced degradation in classification performance when models trained on one acquisition site were applied to data from the other without correction (Fig. 6). This effect was observed across feature representations and model classes, indicating a substantial impact of scanner- and protocol-related domain shift. For FC, direct transfer to IKEM reached 67.2–73.4% balanced accuracy and transfer to NUDZ 63.8–69.9% across the four representative classifiers. This was well above the 50% chance level but 5 to 12 percentage points below the within-site reference (for RR, 73.4% against 78.5% at IKEM and 68.1% against 77.4% at NUDZ).

**Figure 6:**
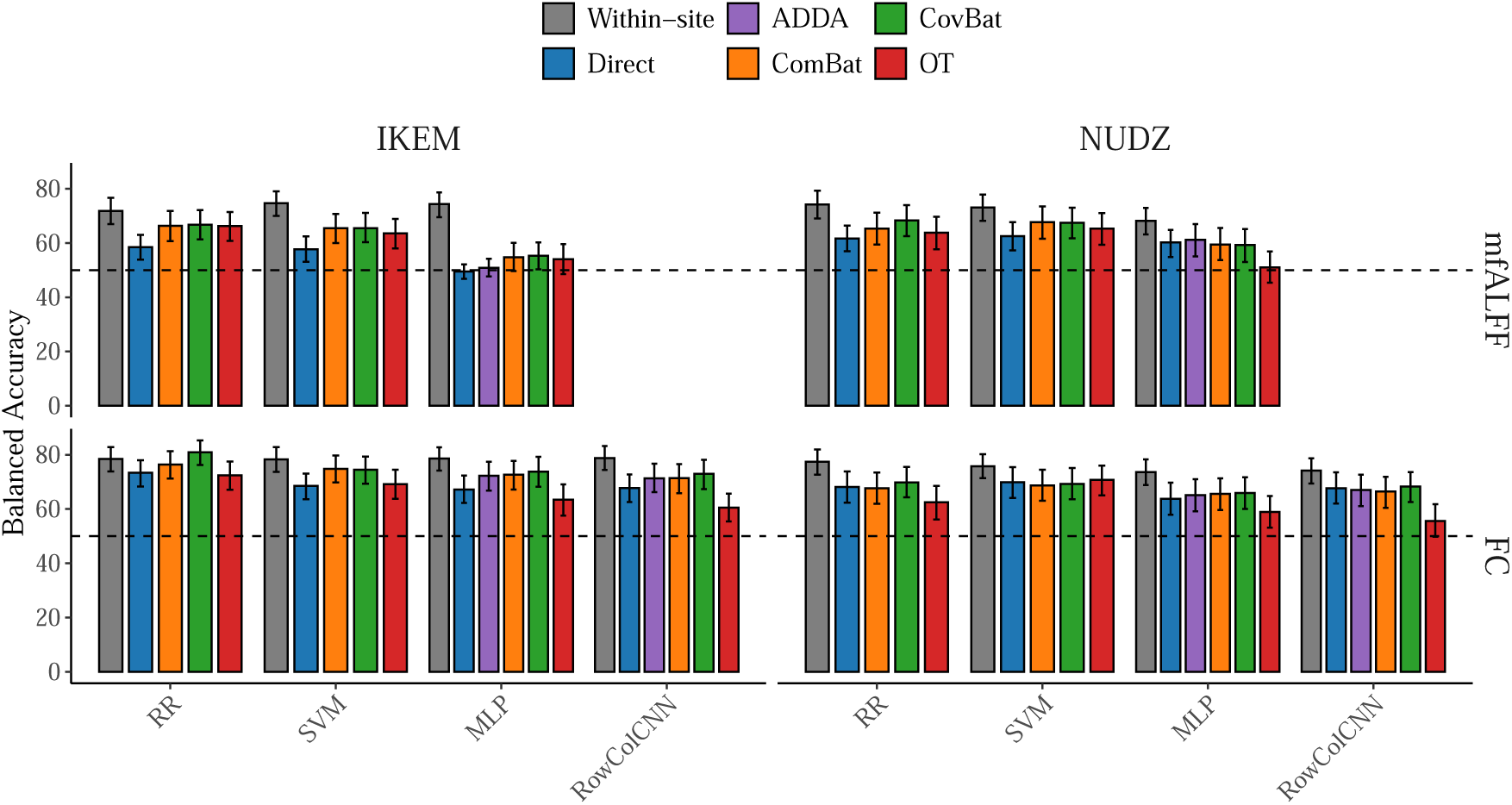
Cross-site classification performance and effect of distribution-alignment strategies. Balanced accuracy (mean *±* 95% bootstrap confidence intervals; 2000 bootstrap resamples) evaluated in the cross-site setting, where models were trained on one acquisition site and evaluated on the other. Results are shown separately for both transfer directions (grid columns, labeled by target site) and for functional connectivity (FC) features and mean-normalized fractional ALFF (mfALFF) features (grid rows), for four representative classifiers (RR, SVM, MLP, and RowColCNN). The RowColCNN operates on connectivity matrices and is therefore absent from the mfALFF row. The horizontal dashed line indicates chance-level performance (50% balanced accuracy). Performance is reported for unaligned models trained directly on the source site (Direct), for feature-level distribution alignment using ComBat, CovBat, and OT, and for latent-level domain adaptation applied to NN models using ADDA. A within-site reference (Within-site), trained and evaluated on the target site, is shown for comparison.

Incorporating distribution-alignment strategies partially mitigated this degradation, but only when transferring to IKEM. Each strategy was tested against its own direct-transfer baseline. For transfer to IKEM, CovBat improved FC accuracy significantly for three of the four classifiers (by 5.9 to 7.6 percentage points) and ComBat for the SVM (by 6.3 points). The remaining harmonization gains, 3.0 to 5.5 points, did not reach significance. Among the deep adaptation methods, ADDA improved MLP transfer significantly (by 5.1 points) and WDGRL by 6.8 points without reaching significance (*p* = 0.074), whereas DANN, DeepCORAL, and DAN gained at most 1.5 points. No deep method improved the RowColCNN significantly (0.1 to 3.6 points). No deep method differed significantly from another or from feature-level harmonization (Supplementary materials). For transfer to NUDZ, no strategy improved accuracy significantly (deltas between *−*2.5 and +3.3 points). No strategy other than OT significantly reduced accuracy for any classifier, feature, or direction. OT changed FC transfer by *−*12.0 to +0.8 points, with significant losses for the RowColCNN in both directions.

The mfALFF feature followed the same direction-dependent pattern. Direct transfer was weaker here (49.5–62.5%). For transfer to IKEM, ComBat and CovBat gained 7.8 to 8.3 points for RR and the SVM, all significantly, and 5.3 to 5.8 points for the MLP without reaching significance. The deep methods gained 1.4 to 16.3 points for the MLP, significantly for all except ADDA, with WDGRL the largest. For transfer to NUDZ, harmonization gained 3.6 to 6.6 points for RR and the SVM without reaching significance, and nothing for the MLP. Unlike on FC, OT also improved mfALFF transfer to IKEM for RR (7.8 points) and the SVM (5.8 points), significantly but not beyond harmonization, whereas it reduced MLP transfer to NUDZ by 9.2 points.

To separate a displaced decision threshold from lost group separation, we compared the operating point of each arm with the ROC curve of direct transfer on the same target (Fig. 7; Supplementary materials). Direct transfer of FC to IKEM displaced the threshold. RR reached a sensitivity of 90.3% at a specificity of 55.6%, and the other three classifiers were more unbalanced still (sensitivity 87.7–93.5%, specificity 37.1–45.2%; Supplementary materials). For RR, ComBat moved the operating point to a sensitivity of 78.6% and a specificity of 74.2% on the direct-transfer ROC curve (offset 0.0 points), with an unchanged AUC. The CovBat operating point was 4.8 points above the curve (*p* = 0.028). The SVM operating points likewise stayed on the curve (offsets 0.6 points for both). For the MLP and the RowColCNN, the aligned operating points lay between 1.3 points below and 3.2 points above the curve, and no strategy changed the AUC significantly (*−*0.5 to +3.2 points). Direct transfer to NUDZ left sensitivity and specificity balanced (66.0% and 70.2% for RR), and the aligned operating points stayed on or near the curve (offsets *−*1.2 to +5.6 points, none significant). In both directions the within-site reference was above the direct-transfer curve (offsets 2.6 to 11.8 points, significant for both neural classifiers at both targets and for RR at NUDZ), and its AUC exceeded that of direct transfer by 4.3 to 10.2 points. OT fell on or below the curve (offsets 0.0 to *−*9.3 points) and lowered the AUC by 3.2 to 11.2 points. The mfALFF feature showed the same pattern (Supplementary materials).

**Figure 7:**
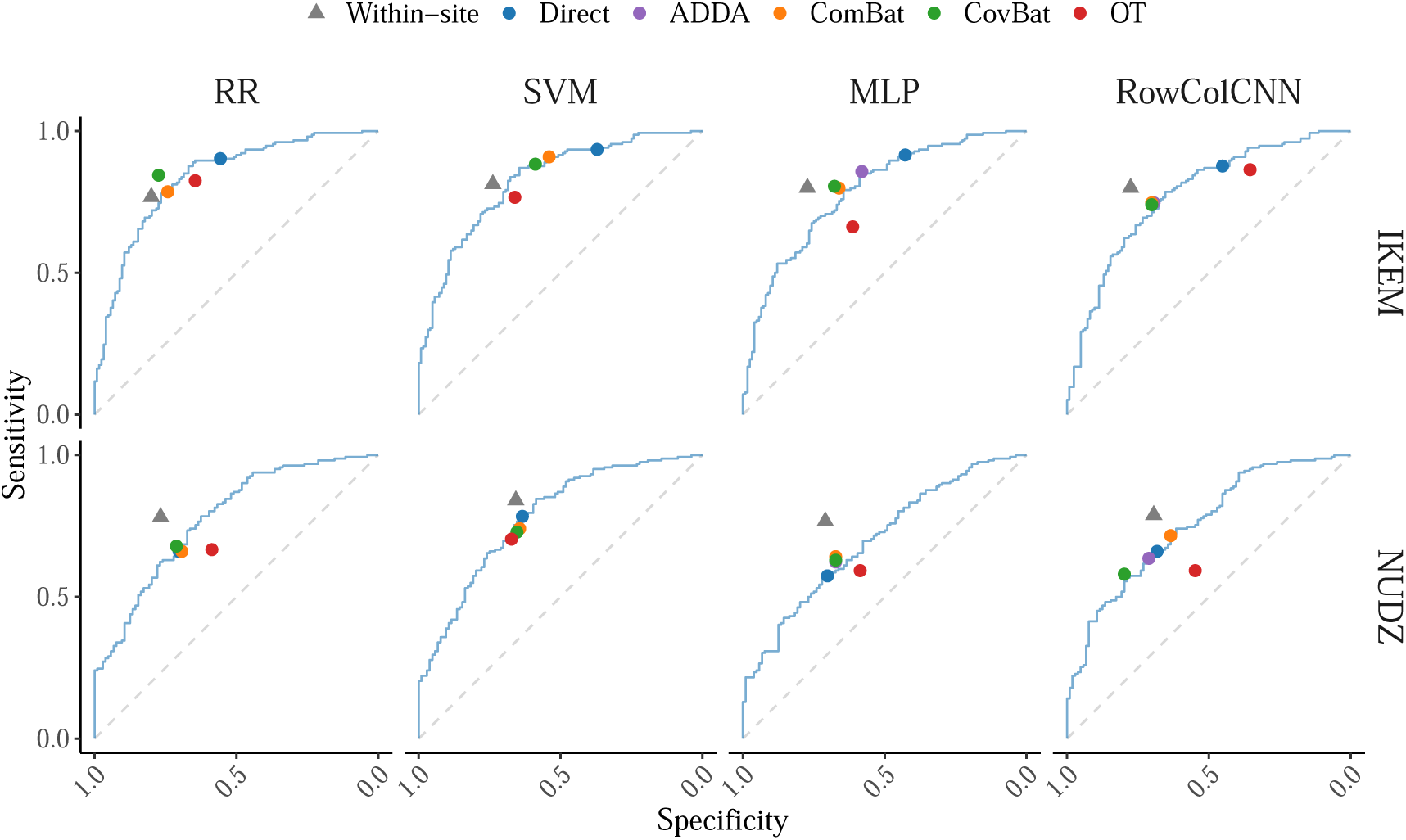
Operating points of the cross-site strategies relative to the ROC curve of direct transfer. Each panel shows the ROC curve of unaligned direct transfer (Direct) for one classifier (columns: RR, SVM, MLP, and RowColCNN) and one target site (rows: IKEM, NUDZ), computed from the averaged class probabilities on functional connectivity (FC) features. Points mark the specificity and sensitivity of each strategy at a probability threshold of 0.5: Direct, ADDA (neural models only), ComBat, CovBat, OT, and the within-site reference (Within-site, triangles). The dashed diagonal marks chance-level performance. For the SVM, probabilities are Platt-scaled, so its operating points need not coincide with its predicted labels.

Taken together, direct transfer cost 5 to 12 points of balanced accuracy. For transfer to IKEM, part of this loss came from a displaced decision threshold, which harmonization and deep adaptation corrected, recovering between about half and all of the gap to the within-site reference. The rest reflected reduced group separation, which alignment did not significantly recover, and OT significantly improved FC transfer in none of the eight comparisons.

## 4 Discussion

This study evaluated TML and NN approaches for classifying individuals with FES using multimodal neuroimaging data acquired at two sites within the same study. Under a rigorously controlled experimental design, we compared unimodal and multimodal representations, assessed performance under matched acquisition conditions, and quantified degradation under cross-site transfer. We further examined whether distribution-alignment strategies could mitigate scanner-related domain shift. Among these findings, the pronounced loss of performance under cross-site transfer has the clearest implications for clinical generalizability.

### 4.1 Functional connectivity as a robust unimodal representation

Across acquisition sites and model classes, FC emerged as the most robust unimodal representation for FES classification (Fig. 3). This advantage was consistent across classifiers, indicating that it was not driven by a specific modeling choice. The mfALFF was a notable second: despite reducing the functional signal to a single value per region, it was broadly comparable to FC at NUDZ, although the gap was larger at IKEM. In contrast, raw ALFF, VBM, and DWI features showed lower and more variable performance.

These findings align with prior work identifying fMRI-derived features, and FC in particular, as among the most discriminative unimodal markers of schizophrenia Porter et al. [2023], Salvador et al. [2019], Shevchenko et al. [2025]. The robustness of FC likely reflects the ability of network-level functional interactions to generalize more reliably across acquisition settings than structural measures.

mfALFF is a strictly univariate summary. It provides one value per region, describing the amplitude of that region’s own low-frequency fluctuations, with no information about how regions co-vary. It is also two orders of magnitude smaller than the connectivity representation (90 values against 4005 edges in AAL90 atlas). A description of this kind nonetheless performed within a few points of FC at one site.

The agreement analysis indicates that regional amplitude is not a compressed restatement of connectivity. Per-subject agreement between the two feature sets barely exceeded chance for all but one classifier-by-site combination, and in almost every classifier and site a majority of the subjects misclassified from FC were correctly classified from mfALFF (Supplementary materials). The two representations appear to access partly distinct discriminative information. mfALFF therefore remains a viable and much smaller alternative to FC, provided its site differences are harmonized before transfer.

### 4.2 Limited benefit of multimodal feature fusion

Despite the appeal of multimodal neuroimaging for capturing complementary aspects of brain pathology, straightforward feature concatenation provided little benefit over FC. Differences between FC-only and multimodal representations were small and heterogeneous, and augmenting FC with diffusion features tended to degrade rather than improve performance.

This observation is consistent with prior reviews reporting mixed or negligible gains from multimodal fusion in schizophrenia classification Porter et al. [2023], Salvador et al. [2019]. A plausible explanation is that additional modalities contribute information that is partly redundant with FC or less stable across acquisition settings, limiting the effectiveness of simple concatenation in datasets of realistic size. The mfALFF result above constrains this explanation for the functional modalities. Complementary information appears to be present at the subject level, yet concatenating the two feature blocks largely did not recover it. FC+mfALFF did not improve significantly on FC alone for any classifier (*−*0.7 to +1.4 points). Concatenation requires a single classifier to weight 4005 correlation coefficients and 90 amplitude values within one shared feature space. In a sample of this size, the regularization required to control the larger block plausibly suppresses the smaller one. VBM and diffusion features, by contrast, classified close to chance on their own and had little to contribute in combination. Decision-level combination, or any scheme that does not force a single classifier to weight all blocks jointly, may be required to exploit complementary information across modalities.

### 4.3 Comparison of model classes under matched acquisition conditions

Under matched acquisition conditions, TML models and NNs achieved broadly comparable performance (Fig. 5). The paired comparison revealed that RR, the SVM, the MLP, and the RowColCNN consistently ranked among the most robust performers, whereas the RF and the more complex graph architectures GCN and GAT showed worse or more variable performance (Supplementary materials).

These results indicate that, in well-controlled within-site settings, model simplicity and effective regularization may be more important than architectural complexity. Regularized linear models were sufficient to extract discriminative information from FC, consistent with prior work Vieira et al. [2020]. Among neural approaches, the competitive performance of the MLP and of the RowColCNN, which treats the connectivity matrix as an image rather than as a graph, suggests that modest nonlinear learning can be beneficial without explicit graph modeling.

### 4.4 Cross-site generalization and the impact of scanner-related domain shift

Cross-site evaluation revealed a pronounced degradation in classification performance when models trained on one site were applied to data from another without correction (Fig. 6). This effect was observed across feature representations and model classes, underscoring the substantial impact of scanner- and protocol-related domain shift even within a single study. As expected, strong within-site performance did not fully translate into cross-site generalization: direct transfer remained well above chance but lost 5 to 12 percentage points of balanced accuracy relative to the within-site reference.

These findings are consistent with prior reports of limited cross-site generalizability in schizophrenia classification Porter et al. [2023], Vieira et al. [2020]. Notably, the degradation occurred despite identical preprocessing pipelines, indicating that scanner-related effects can partly mask disease-related signals and cannot be addressed through model selection alone.

The size of this loss should be interpreted in the context of the cohort design. The two sites belong to a single study: the study design and recruitment were shared, and acquisition was kept as similar as possible, although a newer scanner was later established at NUDZ and participant characteristics differed (Table 1). The remaining between-site variation therefore stems from the change of scanner and its protocol and from the participant samples, rather than from the differences in study design and recruitment that characterize many independently collected multi-center datasets Porter et al. [2023], Vieira et al. [2020].

Within site, the IKEM sample was somewhat easier to classify than NUDZ, by about three percentage points on average for the four leading classifiers on FC (79.3% against 76.2%), although only by one point for RR. The gap is modest and of the same order as the confidence intervals on the individual estimates, so it may simply reflect sampling variability. The clinical variables recorded for the patients do not account for it in the obvious direction. The NUDZ patients had somewhat lower global functioning (mean GAF 64.6 against 70.6) and a longer interval from diagnosis to scanning (9.1 against 6.3 months), yet were the harder group to classify. Greater symptom burden and a longer interval since diagnosis would, if anything, be expected to make the patient group more separable, so the site difference is not explained by clinical severity or illness timing.

### 4.5 Effectiveness of distribution-alignment strategies

Feature-level distribution-alignment strategies partially mitigated cross-site performance degradation when transferring to IKEM, and the deep adaptation methods did so less consistently. CovBat improved FC transfer to IKEM significantly for three of the four classifiers and ComBat for one, and no method other than OT significantly reduced accuracy in either direction. Of the deep methods, only ADDA reached significance on FC, but the methods behaved as one group (Supplementary materials): CovBat was the most consistent, but it differed significantly from ComBat only for RR, and no deep method differed significantly from another or from feature-level harmonization. On mfALFF, the deep methods gave the largest gains, up to 16.3 points for WDGRL.

The effectiveness of feature-level alignment is notable given its simplicity and model-agnostic nature. By operating directly on input feature distributions, these approaches reduce site-related variability prior to classifier training, benefiting both TML and NN models.

The operating-point analysis (Fig. 7) separates two ways in which direct transfer can lose balanced accuracy. It can displace the decision threshold, so that a model calibrated on one site labels too many target subjects as patients or as controls, which lowers balanced accuracy but leaves the AUC unchanged. It can also weaken the separation between the groups, which lowers the AUC as well. Direct transfer to IKEM mainly displaced the threshold, and harmonization corrected it. For the linear classifiers, most aligned operating points moved along the direct-transfer ROC curve toward a balanced sensitivity and specificity, raising balanced accuracy with the AUC almost unchanged. For the neural classifiers, the AUC rose by at most 3.2 points, without reaching significance. For transfer to IKEM, the best strategy raised balanced accuracy by 5.3 to 7.6 points for every classifier. This recovered the whole balanced-accuracy loss for RR, whose loss was the smallest, and about half to two thirds of it for the other classifiers. The within-site operating points nevertheless lay above the direct-transfer ROC curve, with a higher AUC. Alignment of the input or latent distributions did not restore this lost separation. Direct transfer to NUDZ left sensitivity and specificity balanced, so there was little threshold displacement to correct, consistent with the absence of a significant gain in balanced accuracy in that direction.

Optimal-transport alignment, by contrast, did not significantly improve FC transfer for any classifier and reduced it significantly for the RowColCNN. On the one-dimensional decision score of the source classifier, the exact transport plan paired the subjects of the two sites in order of their score. Transport then replaced the full feature vector of each source subject with that of its target partner, or the average of two. The classifier was therefore refit on target data carrying the labels of the source partners, which are correct only where the source classifier already orders the target subjects correctly. These labels add no information beyond direct transfer, and in the 4005-dimensional FC space the classifier plausibly fits their errors. On the 90 mfALFF values the transport did less harm and improved transfer to IKEM for the linear classifiers, but not beyond harmonization.

### 4.6 Methodological implications for neuroimaging-based classification

Most importantly, these results demonstrate that robustness to scanner-related domain shift should be treated as a primary evaluation criterion rather than a secondary validation step. Models that perform well under matched acquisition conditions may lose a substantial part of their accuracy across sites, and explicit alignment strategies recovered only part of this loss.

Because alignment worked mainly by recalibrating the decision threshold, it can be expected to help when direct transfer unbalances sensitivity and specificity, as in transfer to IKEM, but not when they stay balanced, as in transfer to NUDZ. Nor should it be expected to recover the AUC lost in transfer, which fell in both directions.

The present results do not support the view that NNs require substantially more data than a typical neuroimaging study provides to train stably. With nested cross-validation, early stopping, and regularization, the MLP and the FC-specific RowColCNN trained stably on a few hundred subjects, reaching balanced accuracies that were not significantly different from RR and the SVM within site, with overlapping confidence intervals in both transfer directions. Whether larger samples would let them exceed the linear classifiers cannot be determined from our data. At this sample size, their additional capacity found no structure that a regularized linear decision rule had not already captured. The GNNs were the exception. They are the only models that use the connectome as a graph, and they fell significantly below the four leading classifiers at both sites. Whether this reflects the sparsified graph or greater training difficulty is unclear, but modeling the connectome as a graph did not improve classification here. Added complexity should therefore be justified by a demonstrated gain rather than by the expectation that a richer model class will extract more from the same features.

### 4.7 Limitations and future directions

This study was limited to two acquisition sites within a single project, and validation on larger multi-center datasets is required. In particular, the two sites belonged to a single coordinated study but differed in scanner, acquisition protocol, and participant characteristics (Table 1). The direction-dependent benefit of harmonization observed here may not generalize to other site pairs, in which the site shift may displace the decision threshold or reduce group separation to a different degree.

In addition, multimodal integration was restricted to simple feature concatenation. FC and mfALFF misclassified largely different individuals, so complementary information was likely present. Concatenation did not exploit this complementarity, and other fusion strategies were not tested here. The absence of a consistent multimodal gain reported above should therefore be attributed to concatenation rather than to multimodal integration in general.

Future work could extend this framework to prediction of symptom severity, and to systematic investigation of which feature representations remain interpretable and transferable without explicit alignment. Multimodal fusion strategies that model each modality separately, for example by combining the predictions of modality-specific classifiers or by applying a separate penalty to each block, are a natural next step. Such strategies can be judged by whether they recover the subjects that only one of the two modalities classified correctly. Finally, moving beyond diagnostic classification toward prognostic modeling and patient stratification may offer greater clinical relevance and translational impact.

## 5 Conclusion

This study evaluated TML and NN models for FES classification using multimodal neuroimaging data acquired at two sites within the same study. We examined the effect of feature representation, model class, and distribution-alignment strategies under both matched and cross-site acquisition conditions.

Across analyses, FC emerged as the most robust unimodal representation, whereas straightforward multimodal feature fusion did not yield consistent improvements. Under matched acquisition conditions, regularized linear and kernel classifiers performed comparably to the best NNs, whereas the RF and the graph architectures ranked lowest, highlighting the importance of regularization over architectural complexity.

In contrast, direct cross-site transfer led to pronounced performance degradation, underscoring the substantial impact of scanner-related shift. Feature-level distribution-alignment strategies and, less consistently, deep adaptation partially mitigated this degradation in one transfer direction, mainly by correcting a displaced decision threshold, whereas optimal transport did not improve functional connectivity transfer. Alignment is therefore most likely to help when direct transfer unbalances sensitivity and specificity.

Overall, for sites of a single coordinated study, our findings indicate that strong unimodal models combined with principled handling of scanner shift offer a more reliable foundation for generalizable neuroimaging-based classification in FES than focusing on classifier design alone.

## CRediT authorship contribution statement

**Dominik Klepl**: Conceptualization, Formal analysis, Funding acquisition, Investigation, Methodology, Software, Validation, Visualization, Writing – original draft, Writing – review and editing. **Barbora Rehák Bučková**: Conceptualization, Data curation, Investigation, Methodology, Supervision, Writing – review and editing. **Jakub Svoboda**: Conceptualization, Data curation, Investigation, Methodology, Writing – review and editing. **David Tomeček**: Data curation, Writing – review and editing. **Filip Španiel**: Data curation, Funding acquisition, Writing – review and editing. **Jaroslav Hlinka**: Conceptualization, Formal analysis, Funding acquisition, Investigation, Project administration, Resources, Supervision, Validation, Writing – original draft, Writing – review and editing.

## Financial disclosure

This work is supported by CAS program Strategy AV21 "AI" AV21-VP34/2025, ESO trajectory - Czech Health Research Council Project No. NU21-08-00432, PPPLZ - the Czech Academy of Sciences project PPLZ L100302451, and ERDF-Projects BRADY, No. CZ.02.01.01/00/22_008/0004643 and BRAINSCAPE, No. CZ.02.01.01/00/23_020/0008560. Computational resources were provided by the e-INFRA CZ project (ID:90254), supported by the Ministry of Education, Youth and Sports of the Czech Republic.

## Conflict of interest

The authors declare no potential conflicts of interest.

## Data and code availability

The data that support the findings of this study are available from the corresponding author upon reasonable request. The code used for all analyses and experiments is publicly available at https://github.com/dominikklepl/Multimodal-FES-Classification. The repository also contains full configuration details for all trained models, including complete hyperparameter specifications.

## Declaration of generative AI and AI-assisted technologies in the manuscript preparation process

During the preparation of this work, the authors used ChatGPT (OpenAI) and Claude (Anthropic) to assist with language editing, restructuring of text, preparation of LaTeX tables, and writing and optimization of code used for analysis. After using these tools, the authors reviewed and edited all content as needed and take full responsibility for the content of the published article.

## Notes

### Competing Interest Statement

The authors have declared no competing interest.

### Summary of Updates

Revised following peer review: analyses recomputed on uniformly preprocessed data from a larger cohort (544 subjects), additional alignment methods and atlases, updated title, abstract, results and supplementary materials.

