## Supplementary materials for "Domain Adaptation Improves Cross-site Classification of First-episode Schizophrenia from Multimodal Neuroimaging Data"

#### 1 Hyperparameter search spaces

We report the hyperparameter search spaces used during model configuration and optimization. For each model class, we specify the full set of tunable parameters, their sampling distributions, and admissible ranges or categorical options.

Search spaces are provided separately for ridge regression (RR) (Table 1), support vector machine (SVM) (Table 2), and random forest (RF) (Table 3). For neural network (NN) models, we report shared training and classifier-level hyperparameters (Table 4), as well as architecture-specific search spaces for the multilayer perceptron (MLP) (Table 5), row- and column-wise convolutional neural network (RowColCNN) (Table 6), graph convolutional network (GCN) (Table 7), and graph attention network (GAT) (Table 8). Due to the large number of hyperparameters and the combinatorial number of trained models across modalities and acquisition sites, we refer interested readers to the accompanying repository, where all selected hyperparameter values for individual models are reported explicitly.

#### 2 Tabular summary of results

For readers who prefer exact numerical values, this appendix reports the balanced-accuracy results underlying the bar charts in the main Results, each given as the mean with its 95% bootstrap confidence interval. Table 9 reports within-site unimodal performance, Table 11 within-site multimodal feature combinations, Table 13 the within-site model comparison, Table 15 cross-site transfer under each distribution-alignment strategy, and Table 17 the subject-level agreement between functional connectivity (FC) and mean-normalized fractional amplitude of low-frequency fluctuations (mfALFF) predictions. Tables 10, 12, 14, and 16 report the area under the ROC curve, sensitivity, and specificity for the same arms. Tables 18 and 19 report, for each cross-site arm, the offset of its operating point from the direct-transfer ROC curve and its change in the area under the ROC curve, for FC and mfALFF features respectively.

#### 3 Additional critical difference analysis results

The figures in this section present pairwise comparisons that further support the findings reported in the main text. Using per-subject correctness and the stratified paired bootstrap, these analyses provide a complementary, distribution-free comparison of models

Table 1: Hyperparameter search space for ridge regression.

| Hyperparameter | Type | Search range / candidates | Sampling | Notes |
|---|---|---|---|---|
| alpha | categorical | {0.001, 0.01, 0.1, 1, 10, 100, 1000} | categorical | — |

Table 2: Hyperparameter search space for the support vector machine.

| Hyperparameter | Type | Search range / candidates | Sampling | Notes |
|---|---|---|---|---|
| kernel | categorical | {linear, rbf, poly, sigmoid} | categorical | — |
| C | float | [0.0001, 100] | uniform | — |
| gamma | categorical | {scale, auto, 0.0001, 0.001, 0.01, 0.1, 1, 10, 100} | categorical | only if kernel $\in$ {rbf, sigmoid} |
| degree | integer | [1, 4] | integer | only if kernel = poly |

and feature representations, highlighting patterns of relative performance that are robust across subjects. Within-site comparisons are shown for unimodal imaging modalities (Fig. 1), for multimodal feature combinations (Fig. 2), and for model classes (Fig. 3), and a cross-site comparison is shown for the distribution-alignment strategies (Fig. 4), illustrating the consistency of the patterns described in the main text. The operating-point analysis for mfALFF features is shown in Fig. 5 and Table 19.

### 4 Full deep domain-adaptation panel

The main-text figures and table show ADDA as the only deep method. Fig. 6 reports the full panel of deep methods evaluated for the neural models, comprising the adversarial approaches ADDA and domain-adversarial neural network (DANN), the Wasserstein-based Wasserstein distance guided representation learning (WDGRL), and the moment-matching approaches DeepCORAL and deep adaptation network (DAN), alongside unaligned direct transfer, for both FC and mfALFF features. For FC, ADDA improved MLP transfer to Institute of Clinical and Experimental Medicine (IKEM) significantly, by 5.1 points over direct transfer (67.2%), and WDGRL by 6.8 points without reaching significance ($p = 0.074$), whereas DANN, DeepCORAL, and DAN gained 0.8 to 1.5 points. For the RowColCNN, the five methods gained 0.1 to 3.6 points for transfer to IKEM, none significantly. None changed transfer to National Institute of Mental Health (NUDZ) significantly for either classifier ($-2.5$ to $+3.3$ points). On mfALFF the deep methods followed the same direction-dependent pattern as the shallow harmonization methods. For transfer to IKEM they gained 1.4 to 16.3 points for the MLP over direct transfer (49.5%), significantly for all except ADDA, with WDGRL the largest. For transfer to NUDZ they changed accuracy by $-1.3$ to $+2.0$ points.

Table 3: Hyperparameter search space for the random forest classifier.

| Hyperparameter | Type | Search range / candidates | Sampling | Notes |
|---|---|---|---|---|
| n_estimators | integer | [100, 1000] | integer | — |
| criterion | categorical | {gini, entropy, log_loss} | categorical | — |
| max_depth | integer | [5, 50] | integer | — |
| min_samples_split | integer | [2, 20] | integer | — |
| min_samples_leaf | integer | [1, 20] | integer | — |
| min_weight_fraction_leaf | float | [0.0, 0.2] | uniform | — |
| max_features | categorical | {sqrt, log2, None, 0.2, 0.5, 0.8, 1.0} | categorical | — |
| max_leaf_nodes | integer | [10, 100] | integer | — |
| min_impurity_decrease | float | [1e-10, 0.01] | log-uniform | — |

Table 4: Shared training and classifier hyperparameter search space for neural network models.

| Hyperparameter | Type | Search range / candidates | Sampling |
|---|---|---|---|
| learning_rate | categorical | {1e-1, 1e-2, 1e-3, 1e-4, 1e-5} | categorical |
| weight_decay | categorical | {1e-1, 1e-2, 1e-3, 1e-4, 1e-5} | categorical |
| scheduler_factor | float | {0.1, 0.2, . . . , 0.9} | grid (step=0.1) |
| scheduler_patience | integer | [2, 10] | integer |
| classifier_num_layers | integer | [1, 3] | integer |
| classifier_out_channels | categorical | {1, 2} | categorical |
| classifier_dropout | float | {0.00, 0.05, . . . , 0.80} | grid (step=0.05) |

### 5 Atlas robustness

To verify that the main findings are not an artifact of the AAL90 parcellation, all experiments were replicated on three additional atlases spanning a range of resolutions: Harvard–Oxford (112 regions), the Schaefer100 cortical parcellation combined with the Tian S2 subcortical atlas (132 regions), and Brainnetome246 (246 regions). Hyperparameters were not re-optimized for any of these parcellations: the configurations selected on AAL90 were reused unchanged. Because these configurations were obtained from source-site data on a different parcellation, the strict separation between configuration selection and cross-site evaluation is preserved, and no target-site information enters model selection. The following figures aggregate the within- and cross-site results across all four atlases (AAL90 together with the three additional parcellations), colored by atlas, to allow direct visual comparison. Across feature selection, model comparison, and adaptation strategy, the patterns closely mirror the AAL90 findings, supporting the robustness of the conclusions to the choice of cortical parcellation even when hyperparameters are transferred without re-tuning.

Table 5: Encoder hyperparameter search space for the multilayer perceptron.

| Hyperparameter | Type | Search range / candidates | Sampling | Notes |
|---|---|---|---|---|
| num_layers | integer | [1, 4] | integer | — |
| hidden_channels_i | categorical | {32, 64, 128, 256, 512, 1024} | categorical | per layer |
| dropout | float | {0.05, 0.10, . . . , 0.80} | grid (step=0.05) | — |

Table 6: Encoder hyperparameter search space for row- and column-wise convolutional neural network.

| Hyperparameter | Type | Search range / candidates | Sampling | Notes |
|---|---|---|---|---|
| row_kernels | categorical | {8, 16, 32, 64, 128, 256, 516, 1024} | categorical | — |
| col_kernels | categorical | {8, 16, 32, 64, 128, 256, 516, 1024} | categorical | — |
| dropout | float | {0.00, 0.05, . . . , 0.80} | grid (step=0.05) | — |
| filter_value | float | {0.1, 0.2, . . . , 1.0} | grid (step=0.1) | top-percentile edge filter |

#### 5.1 Connectivity structure across atlases

The group-level connectivity contrast between patients and controls was broadly similar across all four parcellations (Fig. 7). Regions were ordered within each atlas by hierarchical clustering of the grand-average connectivity, so that the block structure is visible independently of the arbitrary parcel numbering. The patient-minus-control difference matrices showed broadly similar patterns across resolutions, suggesting that the disorder-related connectivity pattern exploited by the classifiers does not depend strongly on the choice of atlas.

#### 5.2 Feature selection across atlases

FC remained the strongest unimodal representation across all four atlases, with mfALFF again the strongest low-dimensional alternative (Fig. 8). Averaged over classifiers and sites, FC reached 76.1% on AAL90, 76.1% on Harvard–Oxford, 78.7% on Schaefer100+Tian S2, and 77.1% on Brainnetome246, and mfALFF 71.6%, 72.0%, 73.0%, and 72.5% respectively, so neither the ordering of the modalities nor the size of the gap between them depended on the parcellation or its resolution. Augmenting FC with additional modalities produced no systematic gain at any resolution (Fig. 9), reproducing the within-site multimodal result from the main text. Diffusion features were not recomputed for the additional parcellations and are therefore omitted from the cross-atlas view.

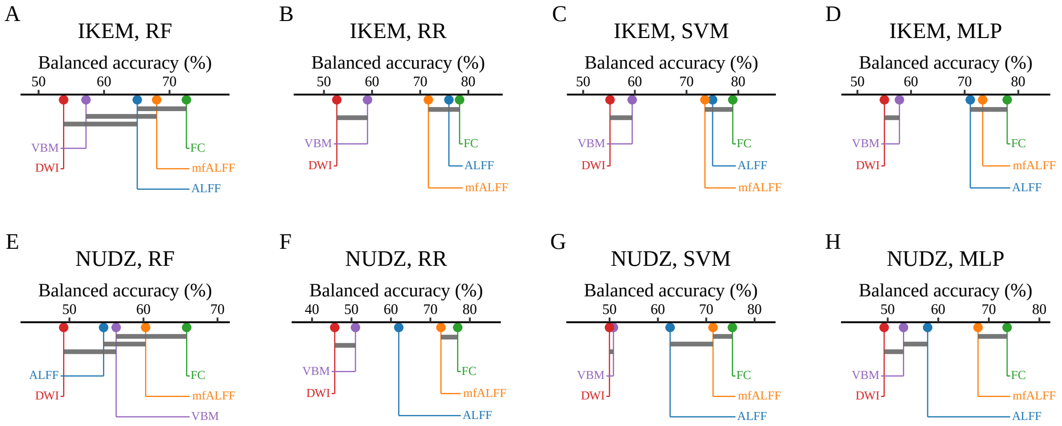


Figure 1: **Within-site comparison of unimodal imaging modalities using critical difference analysis.** Feature representations were scored per subject as correct or incorrect in the within-site setting. Balanced accuracy is shown on the horizontal axis, with higher values indicating better performance. Representations are connected where no pair among them differs significantly under the stratified paired bootstrap with Holm correction over all pairs ($\alpha = 0.05$), indicating no statistically significant difference in performance.

#### 5.3 Model comparison across atlases

Differences between classifiers remained small across atlases (Fig. 10). Regularized linear and kernel classifiers matched or exceeded the neural models at every resolution, and no architecture emerged as systematically superior, reproducing the central model-comparison finding of the main text. The graph neural networks (GNNs) were evaluated on AAL90 only and are therefore omitted from the cross-atlas comparison.

#### 5.4 Distribution alignment across atlases

The cross-site result on FC also held across atlases (Figs. 11 and 12). Averaged over the five classifiers common to all four atlases (RR, SVM, RF, MLP, and the RowColCNN) and over both transfer directions, direct transfer on FC reached 67.7% on AAL90, 70.0% on Harvard–Oxford, 73.0% on Schaefer100+Tian S2, and 69.8% on Brainnetome246, against within-site references of 76.3%, 76.6%, 79.3%, and 77.6% respectively. Direct transfer therefore retained 88 to 92% of the within-site reference at every resolution. ComBat raised these averages by 1.8 to 3.1 points. As on AAL90, the gain was confined to transfer to IKEM. There, ComBat and CovBat improved the SVM significantly at every resolution, CovBat improved the MLP at three of the four, and the RowColCNN improved significantly only with ComBat on Brainnetome246, whereas for RR the gain was significant on AAL90 only. For transfer to NUDZ, no harmonization effect was significant at any resolution, and OT reduced accuracy at every resolution. The area under the ROC curve followed the same pattern (Table 16 for AAL90 and Tables 20, 21, and 22 for the other atlases). Averaged over the five classifiers, direct transfer on FC lowered the AUC by 5.2 to 6.3 points relative to the within-site reference, and ComBat and CovBat changed it by at most 0.8 points at any resolution. For mfALFF, harmonization raised the AUC by 2.1 to 5.4 points on average. The full deep adaptation panel was

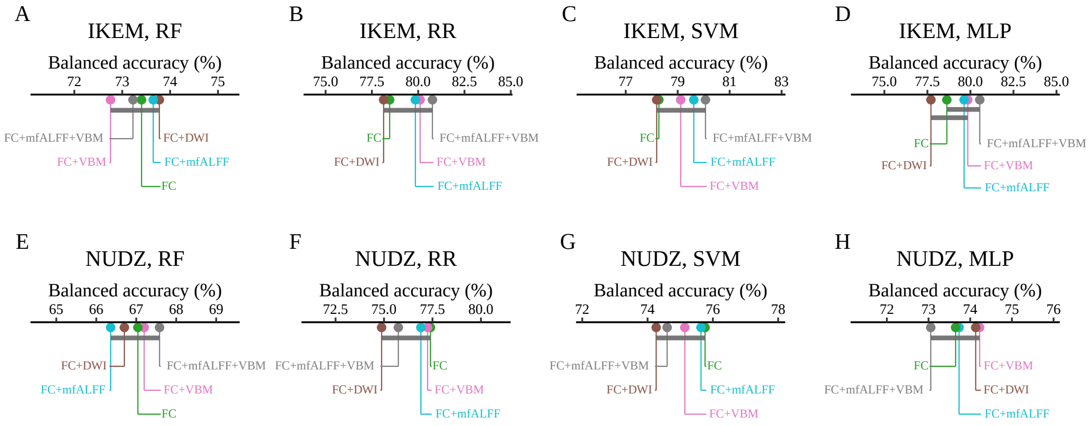


Figure 2: **Within-site comparison of multimodal feature combinations using critical difference analysis.** Feature combinations were scored per subject as correct or incorrect in the within-site setting. Balanced accuracy is shown on the horizontal axis, with higher values indicating better performance. Combinations are connected where no pair among them differs significantly under the stratified paired bootstrap with Holm correction over all pairs ($\alpha = 0.05$), indicating no statistically significant difference in performance. Panels correspond to the RF, RR, SVM, and MLP (columns) at IKEM and NUDZ (rows).

evaluated at every resolution. For the MLP, the deep methods improved transfer to IKEM significantly in 9 of the 20 method-by-atlas comparisons, ranging from all five methods on Schaefer100+Tian S2 to none on Harvard–Oxford, and in one of the 20 for transfer to NUDZ. For the RowColCNN, only one of the 20 comparisons for transfer to IKEM was significant (DANN on Schaefer100+Tian S2, by 7.3 points, $p = 0.017$). For transfer to NUDZ, ADDA lowered accuracy on Brainnetome246 by 7.1 points ($p = 0.002$), the only significant loss from a method other than OT for any classifier, feature set, or atlas.

### References

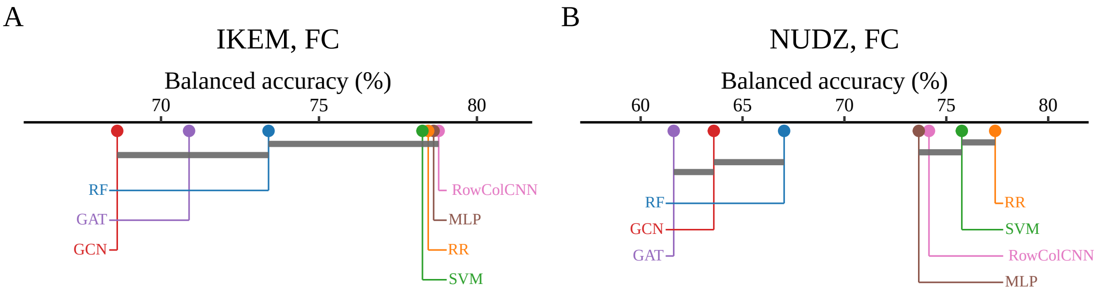


Figure 3: **Within-site comparison of model classes using critical difference analysis.** Diagrams in the style of a critical difference plot summarize pairwise comparisons of model performance in the within-site classification setting, using FC features. Balanced accuracy is shown on the horizontal axis, with higher values indicating better performance. Models are connected where no pair among them differs significantly under the stratified paired bootstrap with Holm correction over all pairs ($\alpha = 0.05$), indicating no statistically significant difference in performance. All evaluated model classes are shown: RR, SVM, RF, MLP, GCN, GAT, and the RowColCNN. The left and right panels correspond to the IKEM and NUDZ cohorts, respectively.

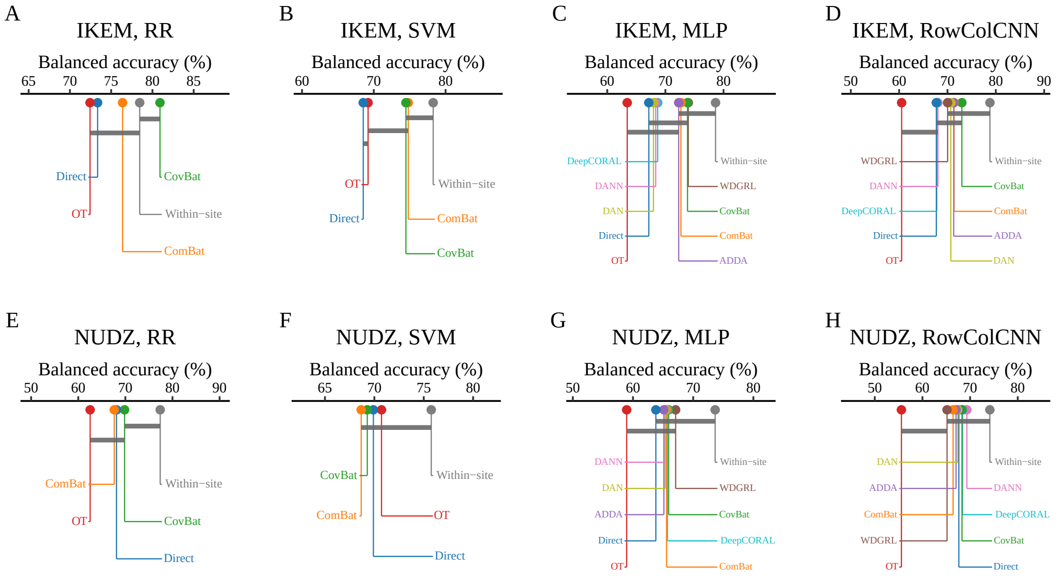


Figure 4: **Cross-site comparison of distribution-alignment strategies using critical difference analysis.** Strategies were scored per subject as correct or incorrect in the cross-site setting on FC features. Balanced accuracy is shown on the horizontal axis, with higher values indicating better performance. Strategies are connected where no pair among them differs significantly under the stratified paired bootstrap with Holm correction over all pairs ($\alpha = 0.05$), indicating no statistically significant difference in performance. Panels correspond to the RR, SVM, MLP, and RowColCNN (columns) transferred to IKEM and to NUDZ (rows). Evaluated strategies include unaligned transfer (Direct), combining batches (ComBat), CovBat, optimal transport (OT), the deep adaptation methods for the neural models, and the within-site reference (Within-site).

Table 7: Encoder hyperparameter search space for the graph convolutional network.

| Hyperparameter | Type | Search range / candidates | Sampling | Notes |
|---|---|---|---|---|
| hidden_channels | categorical | {32, 64, 128, 256, 512, 1024} | categorical | — |
| num_layers | integer | [1, 5] | integer | — |
| dropout | float | {0.05, 0.10, . . . , 0.80} | grid (step=0.05) | — |
| pooling | categorical | {sum, mean, max} | categorical | — |
| jumping_knowledge | categorical | {last, max, cat, none} | categorical | — |
| norm | categorical | {BatchNorm, none} | categorical | — |
| norm_first | categorical | {True, False} | categorical | — |
| dense_first | categorical | {True, False} | categorical | dense projection before message passing |
| linear_first | categorical | {True, False} | categorical | only if dense_first = True |
| linear_hidden | categorical | {32, 64, 128, 256, 512, 1024} | categorical | only if linear_first = True |
| kernels | categorical | {4, 8, 16, 32, 64, 128, 256, 516, 1024} | categorical | only if linear_first = False |
| post_out | categorical | {32, 64, 128, 256, 512, 1024} | categorical | — |
| graph_filter | categorical | {top percentile, KNN} | categorical | — |
| filter_value_p | float | {0.05, 0.10, . . . , 0.50} | grid (step=0.05) | only if graph_filter = top percentile |
| filter_value_k | integer | [1, 30] | integer | only if graph_filter = KNN |
| gradient_clip_value | categorical | {0, 1.0, 2.0} | categorical | — |

Table 8: Encoder hyperparameter search space for the graph attention network.

| Hyperparameter | Type | Search range / candidates | Sampling | Notes |
|---|---|---|---|---|
| hidden_channels | categorical | {32, 64, 128, 256, 512} | categorical | — |
| num_layers | integer | [1, 2] | integer | — |
| v2 | categorical | {True, False} | categorical | GATv2 attention variant |
| dropout | float | {0.05, 0.10, ..., 0.80} | grid (step=0.05) | — |
| pooling | categorical | {sum, mean, max} | categorical | — |
| jumping_knowledge | categorical | {last, max, cat, none} | categorical | — |
| norm | categorical | {BatchNorm, none} | categorical | — |
| norm_first | categorical | {True, False} | categorical | — |
| dense_first | categorical | {True, False} | categorical | dense projection before message passing |
| linear_first | categorical | {True, False} | categorical | only if dense_first = True |
| linear_hidden | categorical | {32, 64, 128, 256, 512, 1024} | categorical | only if linear_first = True |
| kernels | categorical | {4, 8, 16, 32, 64, 128, 256, 516} | categorical | only if linear_first = False |
| post_out | categorical | {32, 64, 128, 256, 512, 1024} | categorical | — |
| graph_filter | categorical | {top percentile, KNN} | categorical | — |
| filter_value_p | float | {0.05, 0.10, ..., 0.50} | grid (step=0.05) | only if graph_filter = top percentile |
| filter_value_k | integer | [1, 30] | integer | only if graph_filter = KNN |
| gradient_clip_value | categorical | {0, 1.0, 2.0} | categorical | — |

Table 9: **Within-site classification performance by unimodal feature.** Balanced accuracy (%, mean with 95% bootstrap confidence interval in brackets) for within-site classification, by imaging modality and classifier, at each acquisition site. The DWI rows are evaluated on the 344 subjects with diffusion data; all other rows use the full cohort. Corresponds to the bar charts in the main Results.

| Modality | Model | IKEM | NUDZ |
|---|---|---|---|
| FC | RR | 78.5 [73.9, 82.8] | 77.4 [72.6, 81.8] |
| FC | SVM | 78.3 [73.8, 82.7] | 75.8 [71.0, 80.4] |
| FC | RF | 73.4 [68.7, 78.0] | 67.0 [62.0, 72.2] |
| FC | MLP | 78.6 [74.2, 82.8] | 73.6 [69.0, 78.2] |
| mfALFF | RR | 71.9 [66.9, 76.5] | 74.2 [69.0, 79.2] |
| mfALFF | SVM | 74.7 [70.0, 79.2] | 73.1 [67.8, 77.8] |
| mfALFF | RF | 69.3 [64.4, 74.5] | 60.1 [55.4, 64.8] |
| mfALFF | MLP | 74.4 [69.6, 78.7] | 68.2 [63.2, 72.8] |
| ALFF | RR | 73.4 [68.8, 78.3] | 61.3 [56.2, 66.4] |
| ALFF | SVM | 73.7 [68.9, 78.5] | 60.1 [55.0, 65.1] |
| ALFF | RF | 62.7 [57.5, 67.8] | 55.0 [50.1, 59.3] |
| ALFF | MLP | 71.4 [66.8, 75.9] | 58.1 [53.7, 62.7] |
| VBM | RR | 58.0 [52.9, 63.0] | 52.9 [49.7, 56.4] |
| VBM | SVM | 57.5 [52.6, 62.5] | 54.1 [50.3, 58.1] |
| VBM | RF | 57.4 [52.0, 62.7] | 55.3 [51.9, 58.9] |
| VBM | MLP | 58.3 [53.6, 63.2] | 54.7 [51.0, 58.4] |
| DWI | RR | 52.7 [46.5, 59.0] | 45.8 [41.3, 50.1] |
| DWI | SVM | 55.2 [48.9, 61.5] | 50.0 [47.2, 52.9] |
| DWI | RF | 53.8 [47.2, 60.7] | 49.2 [43.5, 54.9] |
| DWI | MLP | 55.0 [50.6, 59.5] | 49.3 [45.9, 52.7] |

Table 10: **Within-site unimodal classification: AUC, sensitivity and specificity.** Area under the ROC curve (AUC), sensitivity (Sens.) and specificity (Spec.), in %, each the mean with its 95% bootstrap confidence interval in brackets, for the same arms as Table 9.

| Modality | Model | IKEM | | | NUDZ | | |
|---|---|---|---|---|---|---|---|
| | | AUC | Sens. | Spec. | AUC | Sens. | Spec. |
| FC | RR | 87.8 [83.9, 91.4] | 76.8 [70.6, 82.7] | 80.2 [73.4, 86.3] | 84.2 [79.6, 88.5] | 80.4 [74.8, 85.7] | 74.4 [66.9, 82.1] |
| FC | SVM | 86.7 [82.8, 90.5] | 80.3 [74.4, 85.7] | 76.3 [69.5, 83.0] | 83.7 [79.0, 88.1] | 82.1 [76.8, 87.0] | 69.4 [61.7, 77.1] |
| FC | RF | 81.2 [76.4, 85.9] | 85.2 [80.4, 89.8] | 61.6 [54.2, 69.5] | 78.7 [72.8, 83.6] | 88.9 [84.4, 92.9] | 45.2 [36.9, 54.4] |
| FC | MLP | 86.6 [82.7, 90.2] | 80.0 [74.6, 85.4] | 77.3 [70.8, 83.6] | 80.2 [75.2, 84.8] | 76.9 [70.7, 82.3] | 70.4 [62.5, 78.0] |
| mfALFF | RR | 80.7 [75.6, 85.4] | 76.0 [69.9, 81.7] | 67.7 [60.3, 75.3] | 81.5 [76.2, 86.4] | 81.5 [75.8, 86.3] | 66.9 [58.5, 75.2] |
| mfALFF | SVM | 83.3 [78.5, 87.7] | 80.0 [73.9, 85.4] | 69.4 [62.1, 76.4] | 81.1 [75.9, 85.9] | 80.2 [74.5, 85.2] | 66.0 [57.5, 74.3] |
| mfALFF | RF | 77.5 [72.1, 82.6] | 74.7 [68.0, 80.8] | 63.9 [55.7, 71.8] | 68.4 [62.0, 74.7] | 85.1 [79.8, 89.5] | 35.2 [26.9, 43.5] |
| mfALFF | MLP | 81.3 [76.4, 85.9] | 79.2 [73.3, 84.4] | 69.5 [62.3, 76.8] | 75.1 [69.7, 80.1] | 73.7 [67.8, 79.0] | 62.7 [54.6, 70.6] |
| ALFF | RR | 81.5 [76.3, 86.2] | 78.4 [72.1, 84.1] | 68.4 [61.2, 75.7] | 70.3 [64.5, 75.8] | 74.4 [68.5, 80.1] | 48.1 [39.8, 56.5] |
| ALFF | SVM | 82.6 [77.6, 87.1] | 81.7 [75.8, 87.2] | 65.8 [58.2, 73.0] | 69.1 [63.3, 74.7] | 71.5 [65.2, 77.2] | 48.7 [40.6, 56.8] |
| ALFF | RF | 67.3 [60.8, 73.1] | 78.3 [72.2, 83.9] | 47.1 [38.6, 54.9] | 62.7 [56.1, 69.2] | 81.1 [75.3, 86.1] | 28.8 [21.6, 36.3] |
| ALFF | MLP | 78.4 [73.5, 83.3] | 72.7 [66.4, 78.6] | 70.0 [63.2, 76.3] | 66.6 [60.7, 72.1] | 77.0 [71.4, 82.0] | 39.2 [32.2, 46.7] |
| VBM | RR | 60.2 [53.9, 66.5] | 64.5 [57.7, 70.9] | 51.5 [44.1, 59.0] | 63.3 [56.5, 69.5] | 94.2 [90.8, 97.1] | 11.5 [6.2, 17.2] |
| VBM | SVM | 59.5 [53.6, 65.5] | 64.3 [57.9, 70.4] | 50.6 [43.2, 58.1] | 61.1 [54.3, 67.6] | 89.0 [84.3, 93.0] | 19.2 [12.9, 26.0] |
| VBM | RF | 62.6 [56.1, 69.0] | 69.9 [63.0, 76.5] | 44.8 [36.9, 53.1] | 57.8 [51.2, 64.5] | 93.6 [89.9, 96.9] | 17.1 [10.8, 23.7] |
| VBM | MLP | 61.0 [55.4, 66.5] | 66.4 [60.4, 72.4] | 50.2 [42.8, 57.5] | 58.0 [52.5, 63.3] | 81.6 [77.2, 85.7] | 27.9 [21.9, 33.9] |
| DWI | RR | 54.1 [45.9, 62.3] | 49.2 [40.2, 58.2] | 56.1 [47.2, 65.1] | 46.9 [38.6, 55.0] | 80.0 [73.5, 86.4] | 11.5 [6.0, 18.1] |
| DWI | SVM | 43.0 [38.0, 48.0] | 40.5 [31.5, 49.4] | 69.9 [60.9, 77.8] | 47.1 [41.0, 53.3] | 93.7 [89.5, 97.2] | 6.3 [2.4, 10.9] |
| DWI | RF | 56.0 [47.8, 64.1] | 46.5 [36.4, 56.0] | 61.1 [52.3, 69.8] | 54.7 [46.2, 63.0] | 74.6 [67.0, 82.1] | 23.8 [15.2, 32.8] |
| DWI | MLP | 53.7 [47.8, 59.6] | 42.2 [35.2, 49.1] | 67.8 [62.2, 73.1] | 48.8 [42.4, 55.2] | 84.6 [80.0, 88.8] | 14.0 [9.1, 19.4] |

Table 11: **Within-site classification performance by multimodal feature combination.** Balanced accuracy (%, mean with 95% bootstrap confidence interval in brackets) for within-site classification, by feature combination and classifier, at each acquisition site. Corresponds to the bar charts in the main Results.

| Feature set | Model | IKEM | NUDZ |
|---|---|---|---|
| FC | RR | 78.5 [73.8, 82.6] | 77.4 [72.5, 82.2] |
| FC | SVM | 78.3 [73.7, 82.7] | 75.8 [70.9, 80.5] |
| FC | RF | 73.4 [69.0, 78.1] | 67.0 [62.2, 71.9] |
| FC | MLP | 78.6 [74.1, 82.9] | 73.6 [69.1, 78.4] |
| FC+mfALFF | RR | 79.9 [75.4, 84.1] | 76.9 [71.9, 81.6] |
| FC+mfALFF | SVM | 79.6 [75.2, 84.1] | 75.6 [71.2, 80.3] |
| FC+mfALFF | RF | 73.6 [69.2, 78.2] | 66.4 [61.4, 71.2] |
| FC+mfALFF | MLP | 79.6 [75.4, 84.0] | 73.7 [69.2, 78.3] |
| FC+VBM | RR | 80.1 [75.8, 84.2] | 77.2 [72.4, 81.8] |
| FC+VBM | SVM | 79.1 [74.6, 83.6] | 75.1 [70.3, 79.6] |
| FC+VBM | RF | 72.8 [68.3, 77.2] | 67.2 [62.2, 71.9] |
| FC+VBM | MLP | 79.8 [75.5, 83.8] | 74.2 [69.3, 79.0] |
| FC+DWI | RR | 78.1 [73.6, 82.9] | 74.9 [69.9, 79.5] |
| FC+DWI | SVM | 78.2 [73.7, 82.5] | 74.3 [69.5, 79.2] |
| FC+DWI | RF | 73.8 [69.4, 78.5] | 66.7 [61.7, 71.5] |
| FC+DWI | MLP | 77.7 [73.5, 81.9] | 74.1 [69.5, 78.8] |
| FC+mfALFF+VBM | RR | 80.8 [76.4, 84.8] | 75.7 [71.0, 80.3] |
| FC+mfALFF+VBM | SVM | 80.1 [75.5, 84.3] | 74.6 [69.8, 79.4] |
| FC+mfALFF+VBM | RF | 73.2 [68.7, 77.7] | 67.6 [62.9, 72.4] |
| FC+mfALFF+VBM | MLP | 80.5 [76.4, 84.6] | 73.1 [68.2, 77.6] |

Table 12: **Within-site multimodal classification: AUC, sensitivity and specificity.** Area under the ROC curve (AUC), sensitivity (Sens.) and specificity (Spec.), in %, each the mean with its 95% bootstrap confidence interval in brackets, for the same arms as Table 11.

| Feature set | Model | IKEM | | | NUDZ | | |
|---|---|---|---|---|---|---|---|
| | | AUC | Sens. | Spec. | AUC | Sens. | Spec. |
| FC | RR | 87.8 [83.9, 91.4] | 76.8 [70.6, 82.7] | 80.2 [73.4, 86.3] | 84.2 [79.6, 88.5] | 80.4 [74.8, 85.7] | 74.4 [66.9, 82.1] |
| FC | SVM | 86.7 [82.8, 90.5] | 80.3 [74.4, 85.7] | 76.3 [69.5, 83.0] | 83.7 [79.0, 88.1] | 82.1 [76.8, 87.0] | 69.4 [61.7, 77.1] |
| FC | RF | 81.2 [76.4, 85.9] | 85.2 [80.4, 89.8] | 61.6 [54.2, 69.5] | 78.7 [72.8, 83.6] | 88.9 [84.4, 92.9] | 45.2 [36.9, 54.4] |
| FC | MLP | 86.6 [82.7, 90.2] | 80.0 [74.6, 85.4] | 77.3 [70.8, 83.6] | 80.2 [75.2, 84.8] | 76.9 [70.7, 82.3] | 70.4 [62.5, 78.0] |
| FC+mfALFF | RR | 88.7 [85.0, 91.9] | 78.6 [72.3, 84.4] | 81.1 [74.6, 87.1] | 84.6 [80.0, 89.0] | 81.1 [75.4, 86.2] | 72.7 [64.8, 80.6] |
| FC+mfALFF | SVM | 88.1 [84.4, 91.5] | 81.8 [76.0, 87.0] | 77.4 [70.7, 83.9] | 84.1 [79.3, 88.5] | 83.6 [78.3, 88.3] | 67.7 [59.6, 75.5] |
| FC+mfALFF | RF | 81.9 [77.1, 86.4] | 85.8 [81.2, 90.3] | 61.5 [54.0, 69.0] | 78.5 [72.7, 83.6] | 89.3 [84.7, 93.3] | 43.5 [35.3, 52.7] |
| FC+mfALFF | MLP | 87.2 [83.5, 90.7] | 81.0 [75.5, 86.5] | 78.2 [71.6, 84.3] | 81.6 [76.8, 86.1] | 75.9 [69.9, 81.3] | 71.5 [64.1, 78.8] |
| FC+VBM | RR | 88.5 [84.8, 91.8] | 77.8 [71.5, 83.7] | 82.4 [76.2, 88.2] | 84.2 [79.5, 88.5] | 76.4 [70.5, 82.1] | 78.1 [70.7, 85.4] |
| FC+VBM | SVM | 88.1 [84.2, 91.4] | 80.6 [74.9, 86.2] | 77.6 [70.9, 84.0] | 83.7 [79.0, 88.1] | 82.6 [77.1, 87.5] | 67.7 [59.8, 75.6] |
| FC+VBM | RF | 81.5 [76.7, 86.1] | 84.5 [79.7, 89.4] | 61.0 [53.6, 68.5] | 78.8 [73.0, 83.7] | 89.0 [84.7, 92.9] | 45.4 [37.3, 54.5] |
| FC+VBM | MLP | 87.2 [83.3, 90.7] | 81.9 [76.7, 87.2] | 77.7 [71.5, 83.7] | 80.2 [75.2, 84.8] | 76.9 [71.0, 82.3] | 71.5 [63.8, 79.3] |
| FC+DWI | RR | 86.7 [82.6, 90.5] | 76.1 [69.6, 82.4] | 80.2 [73.6, 86.2] | 82.9 [78.0, 87.5] | 81.5 [76.1, 86.6] | 68.3 [60.2, 76.2] |
| FC+DWI | SVM | 86.6 [82.5, 90.3] | 79.0 [72.7, 84.5] | 77.4 [70.7, 83.7] | 81.8 [76.8, 86.6] | 81.6 [76.2, 86.7] | 66.9 [58.9, 74.7] |
| FC+DWI | RF | 81.7 [77.0, 86.3] | 85.5 [80.7, 90.0] | 62.1 [54.8, 69.7] | 78.4 [72.6, 83.4] | 89.8 [85.4, 93.6] | 43.7 [35.5, 52.9] |
| FC+DWI | MLP | 85.4 [81.3, 89.1] | 79.0 [73.2, 84.5] | 76.5 [69.9, 82.7] | 79.4 [74.6, 84.0] | 75.2 [69.3, 80.6] | 73.1 [65.6, 80.0] |
| FC+mfALFF+VBM | RR | 89.5 [85.9, 92.6] | 78.3 [72.1, 84.2] | 83.2 [77.2, 89.0] | 84.6 [80.1, 88.9] | 83.2 [77.9, 88.1] | 68.3 [60.2, 76.1] |
| FC+mfALFF+VBM | SVM | 88.9 [85.3, 92.2] | 81.4 [75.9, 86.8] | 78.7 [72.1, 85.2] | 84.0 [79.2, 88.3] | 82.5 [77.1, 87.3] | 66.7 [58.8, 74.4] |
| FC+mfALFF+VBM | RF | 82.1 [77.4, 86.7] | 84.7 [79.9, 89.4] | 61.8 [54.4, 69.2] | 78.8 [73.0, 83.7] | 89.0 [84.6, 93.0] | 46.2 [37.9, 55.2] |
| FC+mfALFF+VBM | MLP | 87.1 [83.3, 90.7] | 82.9 [77.6, 88.1] | 78.2 [72.1, 84.2] | 79.1 [74.0, 83.8] | 74.6 [68.6, 80.3] | 71.5 [64.4, 78.9] |

Table 13: **Within-site model comparison.** Balanced accuracy (%, mean with 95% bootstrap confidence interval in brackets) for within-site classification across all seven classifiers on functional connectivity (FC) features, at each acquisition site. Corresponds to the bar charts in the main Results.

| Model | IKEM | NUDZ |
|---|---|---|
| RR | 78.5 [73.8, 82.6] | 77.4 [72.5, 82.2] |
| SVM | 78.3 [73.7, 82.7] | 75.8 [70.9, 80.5] |
| RF | 73.4 [69.0, 78.1] | 67.0 [62.2, 71.9] |
| MLP | 78.6 [74.2, 82.8] | 73.6 [69.0, 78.5] |
| RowColCNN | 78.8 [74.5, 83.1] | 74.1 [69.8, 79.0] |
| GCN | 68.6 [63.8, 73.3] | 63.6 [59.3, 67.9] |
| GAT | 70.9 [66.3, 75.7] | 61.6 [57.4, 65.7] |

Table 14: **Within-site model comparison: AUC, sensitivity and specificity.** Area under the ROC curve (AUC), sensitivity (Sens.) and specificity (Spec.), in %, each the mean with its 95% bootstrap confidence interval in brackets, for the same arms as Table 13.

| Model | IKEM | | | NUDZ | | |
|---|---|---|---|---|---|---|
| | AUC | Sens. | Spec. | AUC | Sens. | Spec. |
| RR | 87.8 [83.9, 91.4] | 76.8 [70.6, 82.7] | 80.2 [73.4, 86.3] | 84.2 [79.6, 88.5] | 80.4 [74.8, 85.7] | 74.4 [66.9, 82.1] |
| SVM | 86.7 [82.8, 90.5] | 80.3 [74.4, 85.7] | 76.3 [69.5, 83.0] | 83.7 [79.0, 88.1] | 82.1 [76.8, 87.0] | 69.4 [61.7, 77.1] |
| RF | 81.2 [76.4, 85.9] | 85.2 [80.4, 89.8] | 61.6 [54.2, 69.5] | 78.7 [72.8, 83.6] | 88.9 [84.4, 92.9] | 45.2 [36.9, 54.4] |
| MLP | 86.6 [82.7, 90.2] | 80.0 [74.6, 85.4] | 77.3 [70.8, 83.6] | 80.2 [75.2, 84.8] | 76.9 [70.7, 82.3] | 70.4 [62.5, 78.0] |
| RowColCNN | 86.9 [83.0, 90.6] | 79.4 [73.6, 84.8] | 78.2 [71.7, 84.3] | 81.3 [76.5, 85.7] | 79.3 [73.8, 84.3] | 69.0 [61.3, 76.7] |
| GCN | 76.7 [71.6, 81.2] | 75.5 [70.0, 80.6] | 61.8 [54.5, 68.9] | 71.1 [65.5, 76.4] | 81.6 [76.8, 86.0] | 45.6 [38.7, 53.0] |
| GAT | 77.5 [72.6, 82.2] | 75.3 [69.6, 80.7] | 66.5 [59.7, 73.0] | 71.8 [66.0, 77.1] | 87.3 [83.2, 91.0] | 36.0 [28.7, 43.6] |

Table 15: **Cross-site transfer: distribution-alignment strategies.** Balanced accuracy (%, mean with 95% bootstrap confidence interval in brackets) for cross-site transfer on functional connectivity (FC) and mean-normalized fractional ALFF (mfALFF) features, by classifier and target site. Within-site is the reference model trained and evaluated on the target site. Remaining columns are unaligned direct transfer (Direct), ComBat and CovBat harmonization, optimal transport (OT), and adversarial discriminative domain adaptation (ADDA). Cells marked n/a are not applicable (ADDA applies only to neural models; RowColCNN operates on connectivity matrices and has no mfALFF variant). Corresponds to the bar charts in the main Results.

| Model | Target | Within-site | Direct | ComBat | CovBat | OT | ADDA |
|---|---|---|---|---|---|---|---|
| *Functional connectivity (FC)* | | | | | | | |
| RR | IKEM | 78.5 [73.8, 82.9] | 73.4 [68.3, 78.0] | 76.4 [71.2, 81.3] | 80.9 [76.3, 85.3] | 72.4 [67.1, 77.5] | n/a |
| RR | NUDZ | 77.4 [72.7, 82.0] | 68.1 [62.4, 73.9] | 67.6 [62.0, 73.5] | 69.8 [64.3, 75.5] | 62.5 [56.2, 68.5] | n/a |
| SVM | IKEM | 78.3 [73.7, 82.8] | 68.5 [63.6, 73.1] | 74.8 [69.8, 79.8] | 74.5 [69.4, 79.3] | 69.2 [63.7, 74.5] | n/a |
| SVM | NUDZ | 75.8 [71.4, 80.2] | 69.9 [64.1, 75.5] | 68.7 [63.0, 74.5] | 69.3 [63.6, 75.1] | 70.7 [65.0, 76.0] | n/a |
| MLP | IKEM | 78.6 [74.1, 82.7] | 67.2 [62.2, 72.4] | 72.7 [67.2, 77.8] | 73.8 [68.2, 79.2] | 63.4 [57.6, 69.1] | 72.3 [66.8, 77.4] |
| MLP | NUDZ | 73.6 [68.9, 78.3] | 63.8 [57.9, 69.7] | 65.6 [59.6, 71.3] | 65.9 [60.1, 71.7] | 59.0 [53.1, 64.8] | 65.1 [59.1, 71.0] |
| RowColCNN | IKEM | 78.8 [74.4, 83.2] | 67.7 [62.6, 72.8] | 71.4 [65.9, 76.6] | 73.0 [67.3, 78.2] | 60.5 [55.5, 65.7] | 71.3 [66.2, 76.7] |
| RowColCNN | NUDZ | 74.1 [69.4, 78.7] | 67.6 [62.0, 73.5] | 66.4 [60.5, 71.9] | 68.3 [62.6, 73.7] | 55.6 [49.8, 61.8] | 67.1 [61.1, 72.7] |
| *mfALFF* | | | | | | | |
| RR | IKEM | 71.9 [67.0, 76.6] | 58.4 [53.9, 63.0] | 66.4 [60.7, 71.8] | 66.7 [61.4, 72.1] | 66.2 [60.8, 71.4] | n/a |
| RR | NUDZ | 74.2 [69.0, 79.3] | 61.7 [57.0, 66.4] | 65.3 [59.4, 71.2] | 68.3 [62.5, 74.0] | 63.8 [57.7, 69.7] | n/a |
| SVM | IKEM | 74.7 [70.0, 79.0] | 57.7 [53.0, 62.5] | 65.5 [60.0, 70.7] | 65.5 [60.3, 71.1] | 63.6 [58.0, 68.8] | n/a |
| SVM | NUDZ | 73.1 [68.2, 77.9] | 62.5 [57.2, 67.7] | 67.7 [61.6, 73.5] | 67.4 [61.7, 73.0] | 65.3 [59.4, 71.0] | n/a |
| MLP | IKEM | 74.4 [69.6, 78.6] | 49.5 [46.8, 52.2] | 54.8 [49.7, 60.1] | 55.3 [50.3, 60.2] | 54.0 [48.5, 59.6] | 50.9 [47.7, 54.2] |
| MLP | NUDZ | 68.2 [63.2, 73.0] | 60.2 [54.8, 64.9] | 59.4 [53.7, 65.6] | 59.3 [53.0, 65.1] | 51.0 [45.3, 56.9] | 61.2 [55.1, 67.0] |

Table 16: **Cross-site transfer: AUC, sensitivity and specificity.** Area under the ROC curve (AUC), sensitivity (Sens.) and specificity (Spec.), in %, each the mean with its 95% bootstrap confidence interval in brackets, for the same arms as Table 15.

| Model | Target | Metric | Within-site | Direct | ComBat | CovBat | OT | ADDA |
|---|---|---|---|---|---|---|---|---|
| *Functional connectivity (FC)* | | | | | | | | |
| RR | IKEM | AUC | 87.8 [83.9, 91.4] | 83.5 [78.7, 88.0] | 83.5 [78.6, 87.9] | 84.2 [79.4, 88.6] | 80.3 [75.1, 85.2] | n/a |
| | | Sens. | 76.8 [70.6, 82.7] | 90.3 [85.5, 94.7] | 78.6 [71.7, 84.8] | 84.4 [78.5, 89.8] | 81.2 [75.0, 87.2] | n/a |
| | | Spec. | 80.2 [73.4, 86.3] | 56.5 [47.9, 65.3] | 74.2 [66.7, 81.9] | 77.4 [70.1, 84.6] | 63.7 [55.4, 72.3] | n/a |
| RR | NUDZ | AUC | 84.2 [79.6, 88.5] | 77.7 [71.9, 82.7] | 77.6 [71.7, 82.7] | 77.7 [71.8, 82.9] | 72.4 [66.1, 78.1] | n/a |
| | | Sens. | 80.4 [74.8, 85.7] | 66.0 [58.9, 73.2] | 66.0 [58.9, 73.2] | 68.5 [61.7, 75.3] | 65.4 [58.1, 72.2] | n/a |
| | | Spec. | 74.4 [66.9, 82.1] | 70.2 [61.3, 78.5] | 69.2 [60.4, 77.6] | 71.2 [62.2, 79.5] | 59.6 [50.4, 68.6] | n/a |
| SVM | IKEM | AUC | 86.7 [82.8, 90.5] | 82.4 [77.6, 86.9] | 82.7 [77.8, 87.2] | 83.3 [78.4, 87.7] | 78.8 [73.6, 83.6] | n/a |
| | | Sens. | 80.3 [74.4, 85.7] | 93.5 [89.6, 97.0] | 88.3 [83.2, 92.9] | 87.7 [82.3, 92.5] | 77.9 [71.2, 84.2] | n/a |
| | | Spec. | 76.3 [69.5, 83.0] | 43.5 [35.2, 52.2] | 61.3 [52.4, 70.1] | 61.3 [52.8, 69.5] | 60.5 [51.8, 68.5] | n/a |
| SVM | NUDZ | AUC | 83.7 [79.0, 88.1] | 78.1 [72.3, 83.2] | 78.0 [72.2, 83.1] | 78.2 [72.4, 83.3] | 73.9 [68.0, 79.8] | n/a |
| | | Sens. | 82.1 [76.8, 87.0] | 73.5 [66.7, 80.1] | 71.0 [64.1, 77.7] | 72.2 [65.6, 78.8] | 72.2 [65.2, 78.7] | n/a |
| | | Spec. | 69.4 [61.7, 77.1] | 66.3 [56.9, 75.5] | 66.3 [56.9, 75.5] | 66.3 [56.9, 75.3] | 69.2 [60.4, 77.9] | n/a |
| MLP | IKEM | AUC | 86.6 [82.7, 90.2] | 78.7 [73.2, 83.5] | 79.1 [73.6, 84.3] | 79.8 [74.3, 84.7] | 67.9 [61.6, 74.1] | 79.7 [74.3, 84.7] |
| | | Sens. | 80.0 [74.6, 85.4] | 91.6 [87.0, 95.5] | 79.2 [72.7, 85.2] | 79.9 [73.2, 85.8] | 65.6 [58.0, 72.9] | 85.7 [80.0, 91.0] |
| | | Spec. | 77.3 [70.8, 83.6] | 42.7 [33.9, 51.2] | 66.1 [57.8, 74.8] | 67.7 [59.5, 76.3] | 61.3 [52.7, 69.6] | 58.9 [50.4, 67.5] |
| MLP | NUDZ | AUC | 80.2 [75.2, 84.8] | 70.1 [63.7, 75.9] | 71.3 [65.3, 77.3] | 71.1 [65.0, 77.2] | 59.7 [52.9, 66.2] | 71.9 [65.7, 77.7] |
| | | Sens. | 76.9 [70.7, 82.3] | 57.4 [49.7, 65.0] | 64.8 [57.6, 72.2] | 63.6 [56.4, 71.0] | 59.3 [51.7, 66.5] | 63.0 [55.8, 70.3] |
| | | Spec. | 70.4 [62.5, 78.0] | 70.2 [61.2, 78.9] | 66.3 [57.4, 75.2] | 68.3 [59.4, 77.2] | 58.7 [49.5, 68.0] | 67.3 [58.2, 76.5] |
| RowColCNN | IKEM | AUC | 86.9 [83.0, 90.6] | 77.7 [72.1, 82.9] | 79.7 [74.2, 84.9] | 79.8 [74.3, 84.8] | 70.0 [64.1, 76.2] | 79.9 [74.4, 84.8] |
| | | Sens. | 79.4 [73.6, 84.8] | 87.0 [81.5, 92.1] | 73.4 [66.2, 80.0] | 73.4 [66.4, 79.9] | 86.4 [81.0, 91.4] | 74.0 [66.4, 80.8] |
| | | Spec. | 78.2 [71.7, 84.3] | 48.4 [39.4, 56.8] | 69.4 [60.8, 76.9] | 72.6 [64.9, 80.5] | 34.7 [26.4, 43.2] | 68.5 [60.2, 76.7] |
| RowColCNN | NUDZ | AUC | 81.3 [76.5, 85.7] | 75.3 [69.2, 80.8] | 75.9 [69.9, 81.4] | 74.8 [68.9, 80.5] | 64.2 [57.4, 70.5] | 75.4 [69.5, 81.0] |
| | | Sens. | 79.3 [73.8, 84.3] | 66.0 [58.7, 72.9] | 70.4 [63.3, 77.1] | 56.8 [49.4, 64.1] | 59.3 [51.2, 66.5] | 63.0 [55.8, 70.5] |
| | | Spec. | 69.0 [61.3, 76.7] | 69.2 [60.2, 77.7] | 62.5 [53.2, 71.7] | 79.8 [72.0, 87.2] | 51.9 [42.6, 62.0] | 71.2 [62.4, 79.6] |
| *mfALFF* | | | | | | | | |
| RR | IKEM | AUC | 80.7 [75.6, 85.4] | 74.1 [68.1, 79.7] | 74.6 [68.4, 80.2] | 74.6 [68.3, 80.1] | 70.8 [64.6, 76.8] | n/a |
| | | Sens. | 76.0 [69.9, 81.7] | 90.3 [85.3, 94.8] | 77.9 [71.1, 84.5] | 78.6 [71.8, 85.2] | 79.2 [72.5, 85.2] | n/a |
| | | Spec. | 67.7 [60.3, 75.3] | 26.6 [19.2, 34.7] | 54.8 [46.0, 63.6] | 54.8 [46.1, 63.9] | 53.2 [44.3, 61.7] | n/a |
| RR | NUDZ | AUC | 81.5 [76.2, 86.4] | 71.3 [65.0, 77.2] | 71.1 [65.0, 77.0] | 72.8 [66.3, 78.7] | 68.3 [61.7, 74.6] | n/a |
| | | Sens. | 81.5 [75.8, 86.3] | 34.0 [26.5, 41.1] | 71.0 [63.6, 78.1] | 72.2 [65.4, 79.1] | 67.9 [60.4, 75.0] | n/a |
| | | Spec. | 66.9 [58.5, 75.2] | 89.4 [83.3, 95.1] | 59.6 [50.0, 69.2] | 64.4 [55.3, 73.9] | 59.6 [50.0, 68.9] | n/a |
| SVM | IKEM | AUC | 83.3 [78.5, 87.7] | 72.7 [66.6, 78.4] | 75.3 [69.1, 80.5] | 74.6 [68.2, 79.9] | 70.5 [64.5, 76.4] | n/a |
| | | Sens. | 80.0 [73.9, 85.4] | 89.6 [84.5, 94.2] | 78.6 [71.9, 85.0] | 78.6 [71.9, 85.1] | 77.9 [71.1, 84.2] | n/a |
| | | Spec. | 69.4 [62.1, 76.4] | 25.8 [18.5, 33.9] | 52.4 [43.6, 61.1] | 52.4 [43.7, 61.2] | 49.2 [40.3, 57.9] | n/a |
| SVM | NUDZ | AUC | 81.1 [75.9, 85.9] | 70.6 [64.1, 76.8] | 72.2 [65.7, 78.1] | 73.6 [67.3, 79.3] | 70.2 [63.5, 76.4] | n/a |
| | | Sens. | 80.2 [74.5, 85.2] | 41.4 [33.5, 48.8] | 71.0 [64.0, 77.8] | 68.5 [61.1, 75.5] | 71.0 [64.0, 77.6] | n/a |
| | | Spec. | 66.0 [57.5, 74.3] | 83.7 [76.1, 90.5] | 64.4 [55.3, 73.8] | 66.3 [56.9, 75.5] | 59.6 [50.0, 69.1] | n/a |
| MLP | IKEM | AUC | 81.3 [76.4, 85.9] | 56.3 [49.6, 63.0] | 60.4 [53.6, 66.8] | 61.2 [54.3, 67.7] | 55.1 [48.4, 61.7] | 57.0 [50.1, 63.7] |
| | | Sens. | 79.2 [73.3, 84.4] | 94.2 [90.3, 97.5] | 80.5 [73.9, 86.4] | 83.1 [76.6, 88.8] | 73.4 [66.4, 79.9] | 92.9 [88.5, 96.7] |
| | | Spec. | 69.5 [62.3, 76.8] | 4.8 [1.6, 8.7] | 29.0 [21.2, 37.5] | 27.4 [19.7, 35.2] | 34.7 [26.7, 42.9] | 8.9 [4.3, 14.6] |
| MLP | NUDZ | AUC | 75.1 [69.7, 80.1] | 62.0 [55.3, 68.5] | 66.6 [59.6, 73.0] | 65.8 [59.0, 72.1] | 52.9 [46.0, 59.6] | 65.0 [58.3, 71.4] |
| | | Sens. | 73.7 [67.8, 79.0] | 35.8 [27.9, 43.0] | 61.1 [53.4, 68.6] | 58.0 [50.6, 65.6] | 65.4 [58.0, 73.0] | 53.1 [45.3, 60.8] |
| | | Spec. | 62.7 [54.6, 70.6] | 84.6 [77.3, 91.2] | 57.7 [48.1, 67.0] | 60.6 [50.8, 69.6] | 36.5 [27.1, 46.5] | 69.2 [60.0, 77.6] |

Table 17: **Subject-level agreement between FC and mfALFF predictions.** Each subject was scored as correct or incorrect under each feature set within site, for the same classifier. Agreement is the percentage of subjects scored alike; chance is the agreement expected from the two marginal accuracies under independence, so it varies by row. The last column is the percentage of the subjects misclassified from FC that were also misclassified from mfALFF.

| Model | Site | Agreement (%) | Chance (%) | $\kappa$ | Shared FC errors (%) |
|---|---|---|---|---|---|
| RR | IKEM | 66.8 | 62.6 | 0.11 | 37 |
| RR | NUDZ | 67.2 | 64.5 | 0.08 | 30 |
| SVM | IKEM | 67.6 | 64.4 | 0.09 | 32 |
| SVM | NUDZ | 67.0 | 63.4 | 0.10 | 33 |
| RF | IKEM | 60.8 | 59.8 | 0.02 | 32 |
| RF | NUDZ | 69.8 | 56.8 | 0.30 | 58 |
| MLP | IKEM | 67.9 | 64.3 | 0.10 | 34 |
| MLP | NUDZ | 65.4 | 59.5 | 0.15 | 42 |

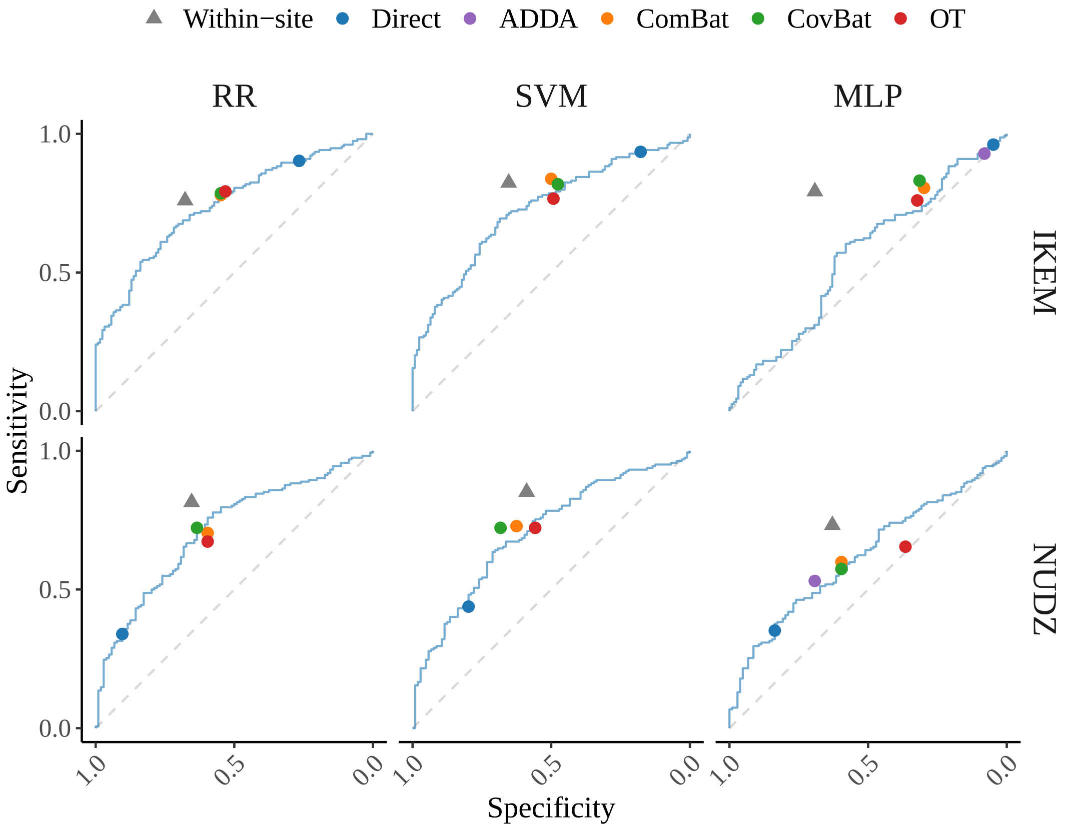


Figure 5: **Operating points of the cross-site strategies relative to the ROC curve of direct transfer for mfALFF features.** Each panel shows the ROC curve of unaligned direct transfer (Direct) for one classifier (columns: RR, SVM, and MLP) and one target site (rows: IKEM, NUDZ), computed from the averaged class probabilities on mean-normalized fractional ALFF (mfALFF) features. Points mark the specificity and sensitivity of each strategy at a probability threshold of 0.5: Direct, adversarial discriminative domain adaptation (ADDA) (MLP only), ComBat, CovBat, OT, and the within-site reference (Within-site, triangles). The dashed diagonal marks chance-level performance.

Table 18: **Operating points against the direct-transfer ROC curve, functional connectivity (FC).** For each arm, the specificity (Spec.) and sensitivity (Sens.) of its pooled class probability at 0.5, in %; the offset, in balanced-accuracy points, from the balanced accuracy of the direct-transfer ROC curve at the same sum of sensitivity and false-positive rate; and the difference in the area under the ROC curve from direct transfer ($\Delta$AUC), each with its 95% bootstrap confidence interval in brackets and paired bootstrap $p$-value (Holm-adjusted over the alignment arms; the within-site arm is not adjusted).

| Model | Target | Arm | Spec. | Sens. | Offset | $p$ | $\Delta$AUC | $p$ |
|---|---|---|---|---|---|---|---|---|
| RR | IKEM | Within-site | 80.0 | 76.8 | 2.6 [-3.3, 8.1] | 0.400 | 4.3 [-0.4, 9.1] | 0.073 |
| | | ComBat | 74.2 | 78.6 | 0.0 [-1.6, 1.3] | 1.000 | -0.0 [-0.6, 0.5] | 0.858 |
| | | CovBat | 77.4 | 84.4 | 4.8 [1.3, 7.8] | 0.028 | 0.7 [-0.5, 2.0] | 0.501 |
| | | OT | 64.5 | 82.5 | -2.6 [-8.1, 2.4] | 0.584 | -3.2 [-7.1, 0.7] | 0.317 |
| RR | NUDZ | Within-site | 76.7 | 78.1 | 8.7 [2.0, 15.2] | 0.010 | 6.5 [0.7, 12.5] | 0.027 |
| | | ComBat | 69.2 | 66.0 | -1.0 [-2.5, 0.0] | 1.000 | -0.1 [-0.4, 0.3] | 1.000 |
| | | CovBat | 71.2 | 67.9 | 1.0 [-1.2, 3.8] | 1.000 | 0.0 [-0.6, 0.7] | 1.000 |
| | | OT | 58.7 | 66.7 | -7.4 [-12.5, -0.6] | 0.118 | -5.3 [-10.1, -0.6] | 0.079 |
| SVM | IKEM | Within-site | 73.9 | 81.3 | 3.8 [-2.4, 9.8] | 0.246 | 4.3 [-0.6, 9.4] | 0.089 |
| | | ComBat | 54.0 | 90.9 | 0.6 [0.0, 1.9] | 1.000 | 0.2 [-0.3, 0.7] | 0.358 |
| | | CovBat | 58.9 | 88.3 | 0.6 [-1.9, 3.2] | 1.000 | 0.8 [-0.1, 1.9] | 0.209 |
| | | OT | 66.1 | 76.6 | -2.6 [-8.9, 3.2] | 1.000 | -3.6 [-7.6, 0.3] | 0.209 |
| SVM | NUDZ | Within-site | 65.8 | 84.1 | 4.3 [-1.6, 11.1] | 0.151 | 5.6 [-0.3, 11.7] | 0.060 |
| | | ComBat | 64.4 | 74.1 | -1.2 [-3.7, 0.6] | 1.000 | -0.1 [-0.6, 0.4] | 1.000 |
| | | CovBat | 65.4 | 72.8 | -1.0 [-3.8, 1.9] | 1.000 | 0.1 [-0.7, 0.8] | 1.000 |
| | | OT | 67.3 | 70.4 | 0.0 [-7.4, 5.6] | 1.000 | -4.2 [-9.4, 0.9] | 0.321 |
| MLP | IKEM | Within-site | 77.3 | 80.0 | 8.5 [2.2, 14.1] | 0.007 | 7.9 [2.8, 13.3] | 0.003 |
| | | ComBat | 66.1 | 79.9 | 1.6 [-0.8, 5.2] | 1.000 | 0.4 [-1.2, 2.0] | 1.000 |
| | | CovBat | 67.7 | 80.5 | 3.2 [0.0, 6.5] | 0.657 | 1.0 [-0.5, 2.7] | 1.000 |
| | | OT | 61.3 | 66.2 | -5.8 [-12.1, -1.3] | 0.178 | -10.8 [-15.2, -6.6] | 0.002 |
| | | ADDA | 58.1 | 85.7 | 0.8 [-1.6, 4.5] | 1.000 | 0.9 [-0.8, 2.8] | 1.000 |
| | | DANN | 52.4 | 85.1 | -1.3 [-4.5, 2.6] | 1.000 | -0.5 [-3.5, 2.4] | 1.000 |
| | | WDGRL | 76.6 | 71.4 | 2.6 [-0.8, 7.1] | 1.000 | 1.5 [-1.5, 4.5] | 1.000 |
| | | DeepCORAL | 50.0 | 88.3 | 0.8 [-3.2, 4.0] | 1.000 | -0.3 [-3.4, 2.8] | 1.000 |
| | | DAN | 48.4 | 88.3 | 0.0 [-3.9, 3.2] | 1.000 | 1.3 [-1.5, 4.2] | 1.000 |
| MLP | NUDZ | Within-site | 71.0 | 76.7 | 11.8 [4.2, 18.4] | 0.002 | 10.2 [2.9, 17.5] | 0.006 |
| | | ComBat | 67.3 | 64.2 | 3.1 [0.0, 6.2] | 0.528 | 1.3 [-1.3, 3.9] | 0.669 |
| | | CovBat | 67.3 | 63.0 | 2.9 [-1.0, 5.8] | 1.000 | 1.0 [-1.8, 3.8] | 0.669 |
| | | OT | 58.7 | 59.3 | -3.7 [-9.6, 1.9] | 1.000 | -10.4 [-16.2, -4.7] | 0.008 |
| | | ADDA | 67.3 | 62.3 | 2.5 [-1.0, 4.9] | 1.000 | 1.8 [-0.9, 4.5] | 0.668 |
| | | DANN | 62.5 | 66.7 | 2.5 [-3.1, 7.4] | 1.000 | 4.2 [0.5, 8.0] | 0.157 |
| | | WDGRL | 65.4 | 69.8 | 5.6 [0.0, 9.9] | 0.483 | 4.2 [0.6, 7.9] | 0.157 |
| | | DeepCORAL | 65.4 | 66.0 | 3.1 [-2.5, 8.6] | 1.000 | 2.9 [-1.0, 6.8] | 0.668 |
| | | DAN | 63.5 | 66.0 | 1.9 [-3.1, 7.7] | 1.000 | 2.8 [-0.9, 6.5] | 0.668 |
| RowColCNN | IKEM | Within-site | 77.6 | 80.0 | 8.3 [2.3, 13.6] | 0.008 | 9.3 [4.0, 14.7] | 0.001 |
| | | ComBat | 70.2 | 74.7 | 1.9 [-2.4, 5.2] | 1.000 | 2.0 [-0.1, 4.2] | 0.383 |
| | | CovBat | 70.2 | 74.0 | 1.3 [-2.6, 4.8] | 1.000 | 2.2 [-0.3, 4.7] | 0.383 |
| | | OT | 35.5 | 86.4 | -4.5 [-8.9, 0.0] | 0.501 | -7.6 [-11.7, -3.6] | 0.002 |
| | | ADDA | 69.4 | 74.7 | 1.6 [-3.2, 5.2] | 1.000 | 2.2 [-0.2, 4.6] | 0.383 |
| | | DANN | 48.4 | 87.7 | 0.6 [-3.9, 5.2] | 1.000 | 3.2 [0.1, 6.4] | 0.295 |
| | | WDGRL | 57.3 | 85.7 | 1.6 [-2.4, 5.8] | 1.000 | 2.5 [-0.4, 5.4] | 0.383 |
| | | DeepCORAL | 49.2 | 87.0 | 0.0 [-3.2, 3.2] | 1.000 | 1.8 [-1.3, 4.8] | 0.383 |
| | | DAN | 52.4 | 88.3 | 1.3 [-2.4, 5.8] | 1.000 | 2.8 [-0.3, 5.9] | 0.383 |
| RowColCNN | NUDZ | Within-site | 69.4 | 78.9 | 7.1 [1.1, 14.2] | 0.021 | 5.9 [-0.4, 12.5] | 0.069 |
| | | ComBat | 63.5 | 71.6 | 0.0 [-3.7, 4.8] | 1.000 | 0.6 [-1.7, 2.9] | 1.000 |
| | | CovBat | 79.8 | 58.0 | 1.2 [-1.9, 5.6] | 1.000 | -0.5 [-2.8, 1.9] | 1.000 |
| | | OT | 54.8 | 59.3 | -9.3 [-15.4, -3.7] | 0.021 | -11.2 [-16.2, -6.3] | 0.002 |
| | | ADDA | 71.2 | 63.6 | 0.0 [-3.7, 3.7] | 1.000 | 0.1 [-1.9, 2.2] | 1.000 |
| | | DANN | 68.3 | 69.1 | 2.5 [-2.9, 6.7] | 1.000 | 0.8 [-2.2, 3.8] | 1.000 |
| | | WDGRL | 61.5 | 70.4 | -1.2 [-5.8, 3.8] | 1.000 | -0.1 [-3.2, 2.8] | 1.000 |
| | | DeepCORAL | 67.3 | 69.1 | 1.9 [-3.1, 6.2] | 1.000 | 0.6 [-2.5, 3.7] | 1.000 |
| | | DAN | 66.3 | 69.1 | 1.9 [-3.8, 5.8] | 1.000 | -0.3 [-3.3, 2.7] | 1.000 |

Table 19: **Operating points against the direct-transfer ROC curve, mfALFF.** For each arm, the specificity (Spec.) and sensitivity (Sens.) of its pooled class probability at 0.5, in %; the offset, in balanced-accuracy points, from the balanced accuracy of the direct-transfer ROC curve at the same sum of sensitivity and false-positive rate; and the difference in the area under the ROC curve from direct transfer (ΔAUC), each with its 95% bootstrap confidence interval in brackets and paired bootstrap $p$-value (Holm-adjusted over the alignment arms; the within-site arm is not adjusted).

| Model | Target | Arm | Spec. | Sens. | Offset | $p$ | ΔAUC | $p$ |
|---|---|---|---|---|---|---|---|---|
| RR | IKEM | Within-site | 67.7 | 76.0 | 4.6 [-1.5, 11.0] | 0.148 | 6.6 [0.3, 12.9] | 0.039 |
| | | ComBat | 54.8 | 77.9 | 0.0 [-4.5, 5.6] | 1.000 | 0.5 [-3.5, 4.6] | 1.000 |
| | | CovBat | 54.8 | 78.6 | 0.0 [-4.0, 6.5] | 1.000 | 0.5 [-3.7, 4.7] | 1.000 |
| | | OT | 53.2 | 79.2 | 0.6 [-3.2, 4.8] | 1.000 | -3.3 [-6.6, -0.0] | 0.149 |
| RR | NUDZ | Within-site | 65.4 | 81.5 | 6.3 [-0.2, 13.4] | 0.055 | 10.1 [3.7, 16.5] | 0.002 |
| | | ComBat | 59.6 | 70.4 | -1.2 [-4.8, 1.0] | 0.899 | -0.2 [-1.7, 1.3] | 0.788 |
| | | CovBat | 63.5 | 72.2 | 0.6 [-2.9, 4.8] | 0.899 | 1.4 [-0.8, 3.7] | 0.439 |
| | | OT | 59.6 | 67.3 | -3.8 [-8.7, 1.9] | 0.899 | -3.0 [-6.5, 0.4] | 0.246 |
| SVM | IKEM | Within-site | 65.3 | 82.3 | 7.4 [1.4, 14.0] | 0.016 | 10.6 [4.4, 17.2] | <0.001 |
| | | ComBat | 50.0 | 83.8 | 3.9 [-1.3, 8.1] | 0.617 | 2.6 [-1.5, 6.9] | 0.438 |
| | | CovBat | 47.6 | 81.8 | 1.9 [-4.0, 6.5] | 1.000 | 1.9 [-2.8, 6.7] | 0.438 |
| | | OT | 49.2 | 76.6 | -1.6 [-5.6, 2.4] | 1.000 | -2.2 [-4.9, 0.6] | 0.389 |
| SVM | NUDZ | Within-site | 58.8 | 85.2 | 7.5 [1.4, 14.8] | 0.018 | 10.5 [3.5, 17.5] | 0.003 |
| | | ComBat | 62.5 | 72.8 | 3.1 [-0.6, 6.7] | 0.309 | 1.6 [-0.4, 3.6] | 0.250 |
| | | CovBat | 68.3 | 72.2 | 4.9 [0.0, 9.3] | 0.147 | 2.9 [0.4, 5.6] | 0.064 |
| | | OT | 55.8 | 72.2 | -1.0 [-5.6, 3.7] | 0.782 | -0.5 [-3.2, 2.2] | 0.732 |
| MLP | IKEM | Within-site | 69.2 | 79.2 | 17.3 [10.1, 23.9] | <0.001 | 25.0 [17.1, 33.1] | <0.001 |
| | | ComBat | 29.8 | 80.5 | 3.9 [-2.4, 8.9] | 0.820 | 4.2 [-2.7, 11.0] | 0.693 |
| | | CovBat | 31.5 | 83.1 | 5.6 [0.0, 10.5] | 0.227 | 4.9 [-2.0, 11.8] | 0.643 |
| | | OT | 32.3 | 76.0 | 1.9 [-3.9, 7.8] | 1.000 | -1.2 [-8.0, 5.7] | 1.000 |
| | | ADDA | 8.1 | 92.9 | -0.8 [-4.0, 2.6] | 1.000 | 0.7 [-4.7, 6.3] | 1.000 |
| | | DANN | 31.5 | 87.0 | 7.3 [1.6, 12.9] | 0.116 | 9.9 [2.7, 17.0] | 0.034 |
| | | WDGRL | 69.4 | 63.6 | 7.8 [0.6, 17.7] | 0.224 | 12.1 [2.9, 21.3] | 0.042 |
| | | DeepCORAL | 35.5 | 86.4 | 9.7 [3.2, 14.5] | 0.024 | 11.1 [3.9, 18.2] | 0.026 |
| | | DAN | 33.9 | 87.7 | 8.9 [3.2, 14.3] | 0.028 | 10.4 [3.5, 17.2] | 0.028 |
| MLP | NUDZ | Within-site | 62.9 | 73.2 | 10.6 [2.9, 18.1] | 0.007 | 13.1 [4.7, 21.1] | 0.002 |
| | | ComBat | 59.6 | 59.9 | 1.9 [-3.1, 6.8] | 1.000 | 4.6 [0.7, 8.6] | 0.146 |
| | | CovBat | 59.6 | 57.4 | 1.0 [-4.3, 5.8] | 1.000 | 3.8 [-0.1, 7.8] | 0.272 |
| | | OT | 36.5 | 65.4 | -7.4 [-12.5, -1.0] | 0.301 | -9.1 [-14.3, -4.0] | 0.005 |
| | | ADDA | 69.2 | 53.1 | 1.9 [-1.2, 6.8] | 1.000 | 3.0 [-0.5, 6.5] | 0.398 |
| | | DANN | 76.9 | 46.3 | 1.0 [-2.9, 5.8] | 1.000 | -0.2 [-3.9, 3.5] | 1.000 |
| | | WDGRL | 76.9 | 47.5 | 1.2 [-2.9, 6.8] | 1.000 | 5.3 [1.1, 9.5] | 0.094 |
| | | DeepCORAL | 71.2 | 45.7 | -1.2 [-5.8, 3.7] | 1.000 | 0.1 [-4.0, 4.2] | 1.000 |
| | | DAN | 75.0 | 44.4 | -1.0 [-4.9, 4.8] | 1.000 | 0.4 [-3.7, 4.5] | 1.000 |

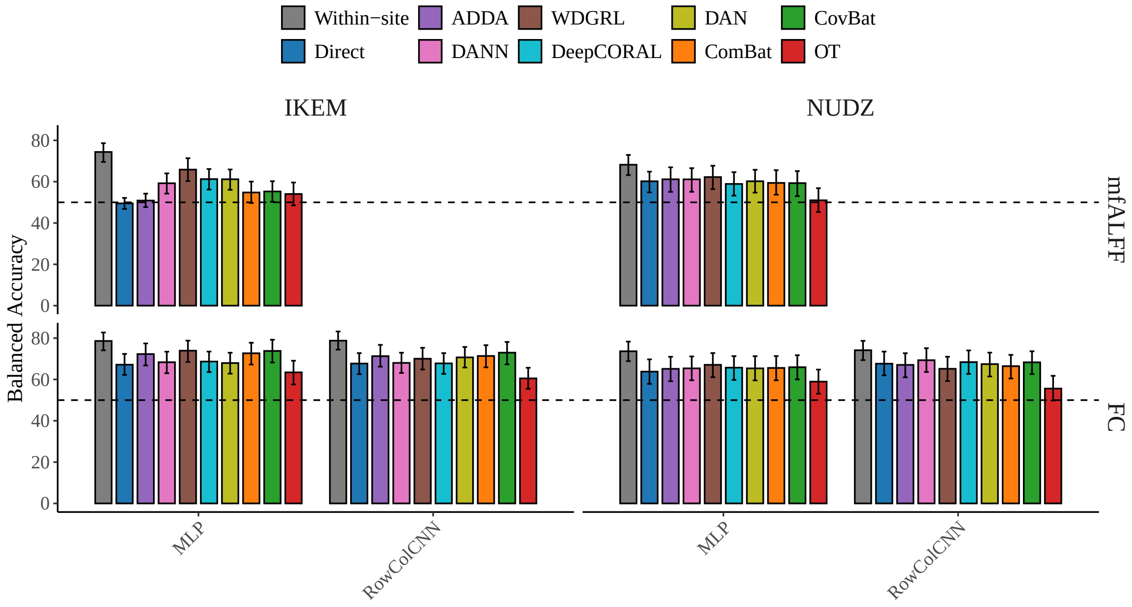


Figure 6: **Cross-site transfer under the full deep domain-adaptation panel.** Balanced accuracy (mean with 95% bootstrap confidence interval; 2000 resamples) for cross-site transfer on functional connectivity (FC) and mfALFF features, shown for the neural models (MLP and RowColCNN; RowColCNN has no mfALFF variant). Strategies comprise unaligned direct transfer (Direct) and the deep adaptation methods ADDA, DANN, WDGRL, DeepCORAL, and DAN. Results are shown separately for both target sites (IKEM and NUDZ).

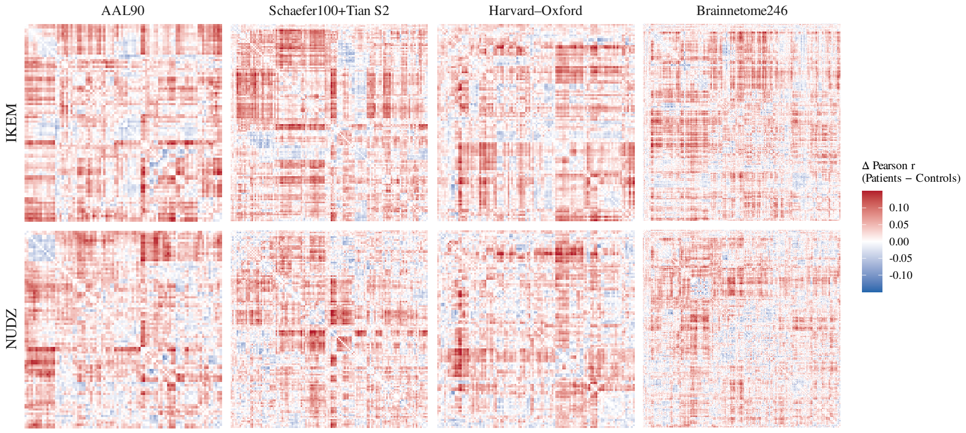


Figure 7: **Patient-minus-control functional connectivity across atlases and sites.** Each panel shows the difference between the mean patient and mean control connectivity matrix (Pearson $r$) for one atlas (columns) and one acquisition site (rows: IKEM, NUDZ). Regions are ordered within each atlas by hierarchical clustering of the grand-average connectivity. Red indicates stronger connectivity in patients, blue stronger connectivity in controls.

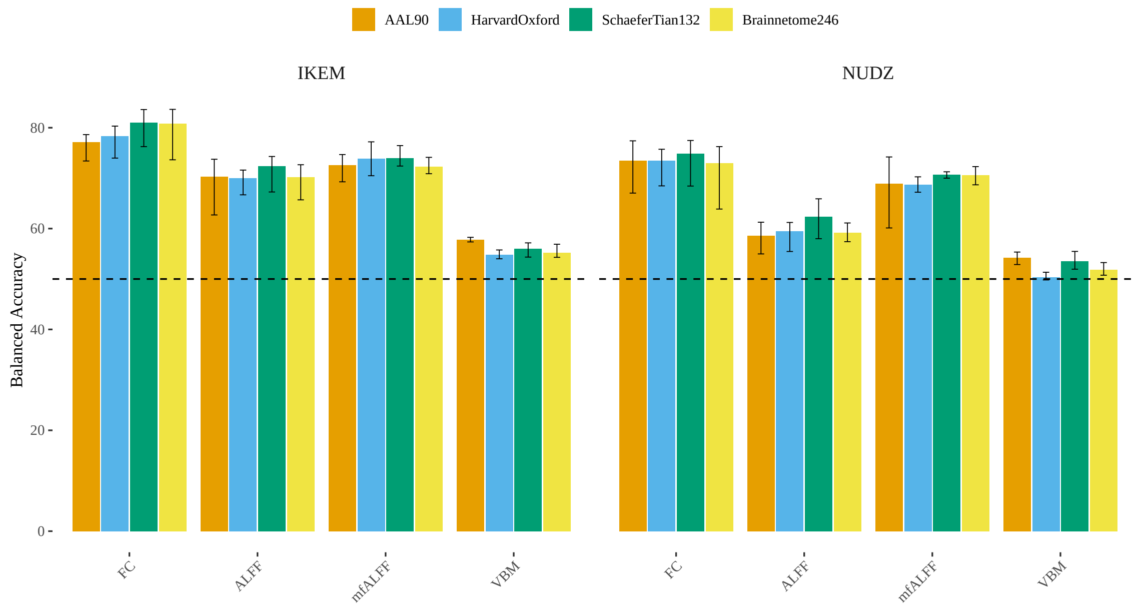


Figure 8: **Within-site unimodal performance across atlases.** Balanced accuracy for within-site classification by unimodal feature, averaged over classifiers and colored by atlas, for both acquisition sites (IKEM, NUDZ). Whiskers span the range across classifiers. Diffusion features are defined on JHU tracts and were joined to the AAL90 export only, so the DWI condition is omitted here.

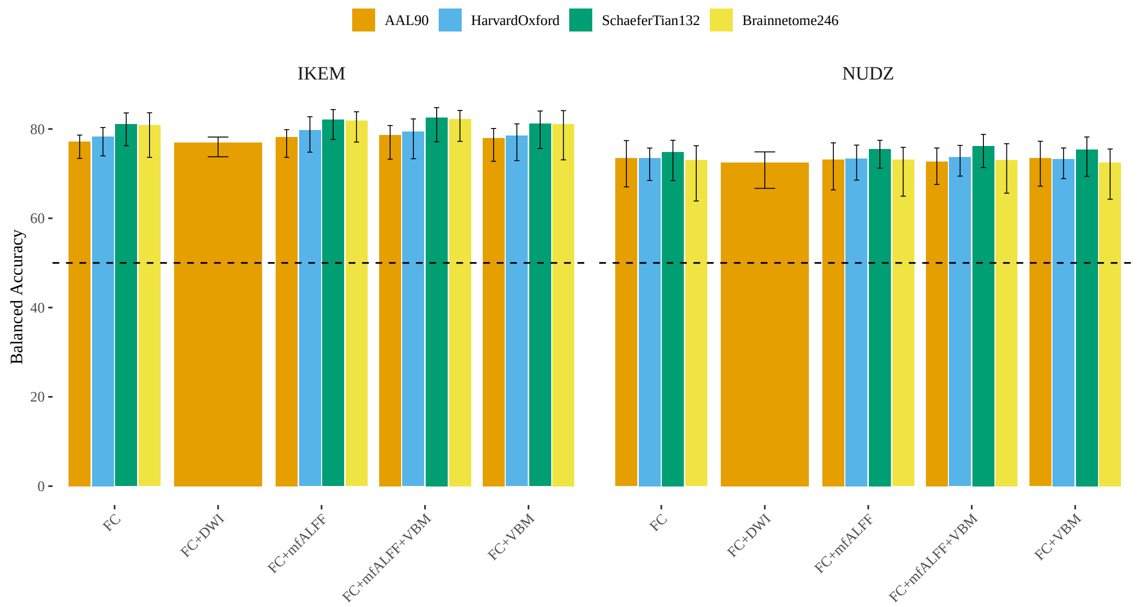


Figure 9: **Within-site multimodal performance across atlases.** Balanced accuracy for within-site classification by feature combination, averaged over classifiers and colored by atlas, for both acquisition sites (IKEM, NUDZ). Whiskers span the range across classifiers. FC+DWI is shown for AAL90 alone, since the diffusion features were joined to that export only.

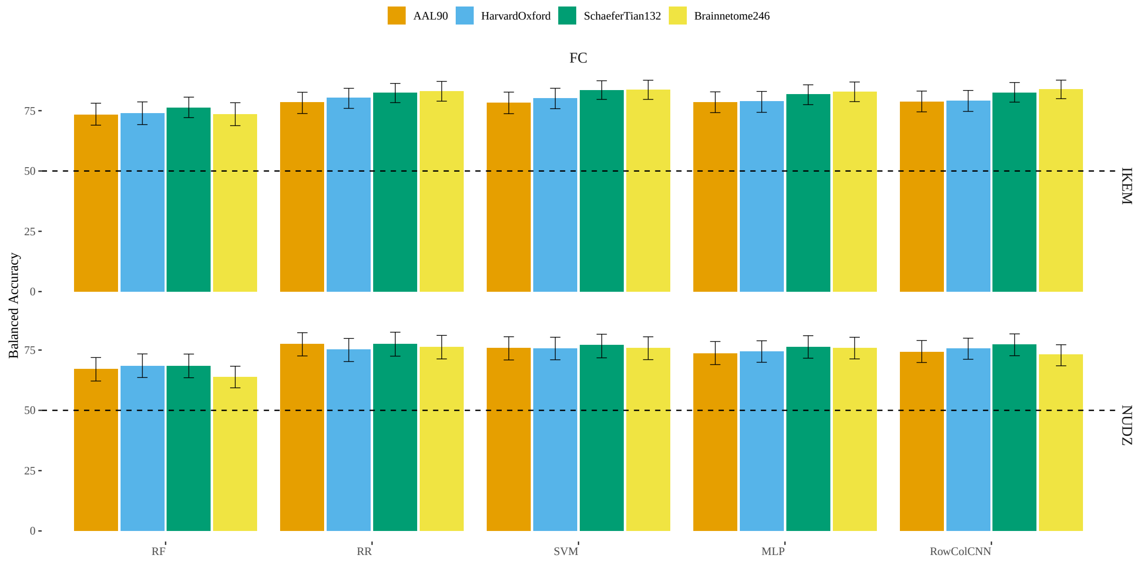


Figure 10: **Within-site model comparison across atlases.** Balanced accuracy for within-site classification on functional connectivity features by classifier, colored by atlas, for both acquisition sites (IKEM, NUDZ). Error bars are 95% bootstrap confidence intervals. The graph neural networks (GCN, GAT) were evaluated on AAL90 only and are not shown.

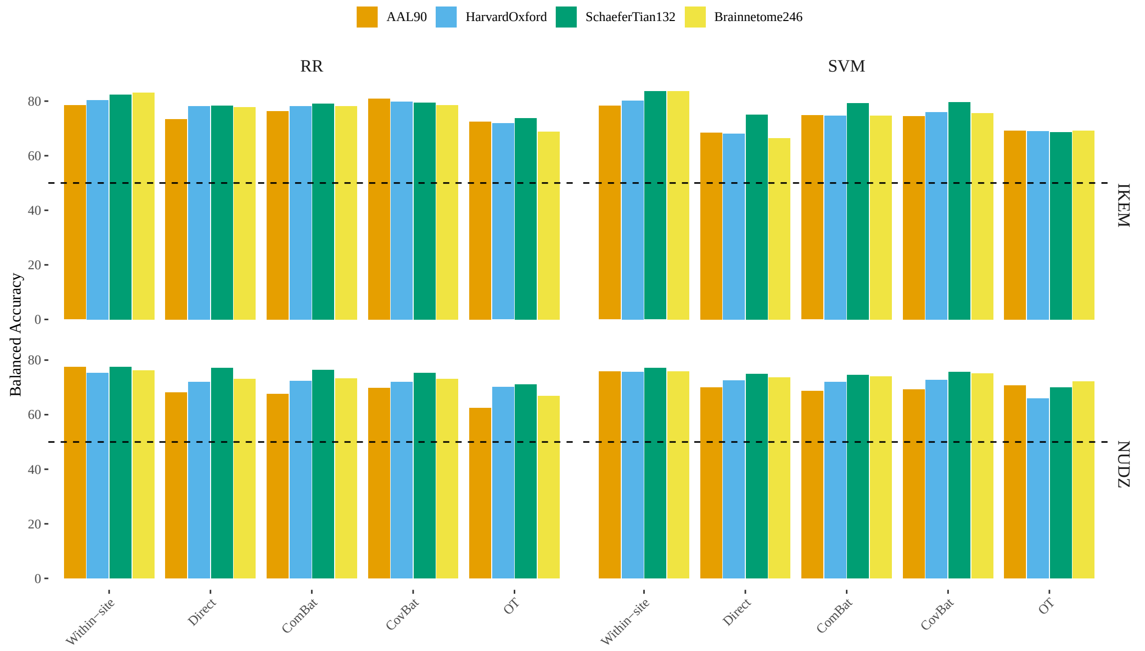


Figure 11: **Cross-site distribution alignment across atlases (traditional machine-learning models).** Balanced accuracy for cross-site transfer on functional connectivity features by alignment strategy, averaged over the traditional machine-learning classifiers and colored by atlas, for both target sites (IKEM, NUDZ). Strategies are the within-site reference (Within-site), unaligned direct transfer (Direct), ComBat and CovBat harmonization, and optimal transport (OT). Bars are means across classifiers and carry no confidence intervals.

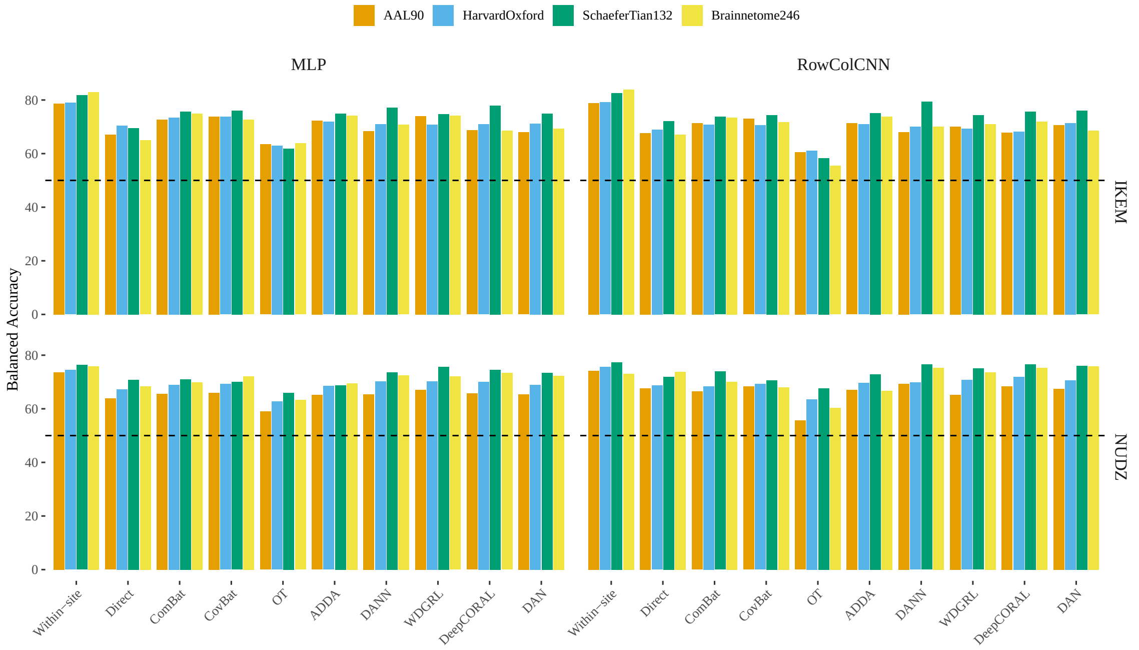


Figure 12: **Cross-site distribution alignment across atlases (neural models).** Balanced accuracy for cross-site transfer on functional connectivity features by alignment strategy, shown separately for the MLP and the RowColCNN and colored by atlas, for both target sites (IKEM, NUDZ). Strategies are the within-site reference (Within-site), unaligned direct transfer (Direct), ComBat and CovBat harmonization, optimal transport (OT), and the deep adaptation methods ADDA, DANN, WDGRL, DeepCORAL, and DAN. The graph neural networks were evaluated on AAL90 only and are omitted.

Table 20: **Cross-site transfer on Harvard–Oxford: AUC, sensitivity and specificity.** Area under the ROC curve (AUC), sensitivity (Sens.) and specificity (Spec.), in %, each the mean with its 95% bootstrap confidence interval in brackets, for cross-site transfer on functional connectivity (FC) and mfALFF features with the Harvard–Oxford atlas, by classifier and target site. Arms as in Table 16.

| Model | Target | Metric | Within-site | Direct | ComBat | CovBat | OT | ADDA |
|---|---|---|---|---|---|---|---|---|
| *Functional connectivity (FC)* | | | | | | | | |
| RR | IKEM | AUC | 89.6 [85.9, 93.1] | 84.5 [79.8, 89.0] | 84.1 [79.2, 88.5] | 84.7 [80.2, 89.1] | 78.2 [72.9, 83.5] | n/a |
| | | Sens. | 79.4 [73.5, 85.0] | 83.1 [77.1, 88.9] | 79.9 [73.3, 86.2] | 80.5 [74.0, 86.7] | 79.2 [72.7, 85.3] | n/a |
| | | Spec. | 81.3 [74.9, 87.1] | 73.4 [65.6, 80.5] | 76.6 [69.0, 83.5] | 79.0 [71.5, 85.8] | 64.5 [55.8, 72.8] | n/a |
| RR | NUDZ | AUC | 85.2 [80.7, 89.2] | 80.0 [74.8, 85.2] | 79.9 [74.7, 85.1] | 80.0 [74.7, 85.2] | 76.2 [70.3, 81.8] | n/a |
| | | Sens. | 77.2 [70.9, 82.7] | 67.9 [61.1, 75.2] | 69.8 [62.8, 76.8] | 71.0 [64.3, 77.8] | 72.8 [66.2, 79.8] | n/a |
| | | Spec. | 73.3 [65.0, 80.8] | 76.0 [67.6, 84.1] | 75.0 [66.7, 83.2] | 73.1 [64.5, 81.5] | 67.3 [58.3, 76.2] | n/a |
| SVM | IKEM | AUC | 89.4 [85.6, 92.8] | 82.7 [77.8, 87.4] | 82.6 [77.7, 87.3] | 83.6 [78.9, 88.2] | 76.1 [70.6, 81.4] | n/a |
| | | Sens. | 83.5 [78.0, 88.6] | 90.3 [85.5, 94.6] | 85.7 [80.0, 91.0] | 85.7 [80.0, 91.1] | 79.2 [73.2, 85.6] | n/a |
| | | Spec. | 76.8 [70.1, 83.2] | 46.0 [37.2, 54.5] | 63.7 [54.7, 71.9] | 66.1 [57.5, 74.2] | 58.9 [50.4, 67.2] | n/a |
| SVM | NUDZ | AUC | 85.4 [80.9, 89.5] | 79.5 [74.1, 84.8] | 79.4 [74.1, 84.7] | 79.7 [74.4, 85.0] | 74.4 [68.5, 79.9] | n/a |
| | | Sens. | 82.2 [76.6, 87.1] | 72.8 [66.3, 79.5] | 72.8 [66.0, 79.4] | 73.5 [67.0, 80.1] | 70.4 [63.4, 77.4] | n/a |
| | | Spec. | 69.2 [61.2, 77.0] | 72.1 [63.5, 80.4] | 71.2 [62.4, 79.4] | 72.1 [63.5, 80.4] | 61.5 [52.5, 70.8] | n/a |
| MLP | IKEM | AUC | 87.2 [83.2, 90.7] | 77.1 [71.4, 82.5] | 78.9 [73.4, 84.2] | 78.4 [72.7, 83.7] | 64.6 [58.0, 71.0] | 79.5 [74.0, 84.6] |
| | | Sens. | 81.3 [75.8, 86.8] | 87.7 [82.5, 92.4] | 79.2 [72.7, 85.2] | 80.5 [74.2, 86.5] | 66.2 [58.7, 73.6] | 82.5 [76.4, 88.1] |
| | | Spec. | 76.6 [69.8, 83.0] | 53.2 [44.2, 61.5] | 67.7 [58.9, 75.6] | 66.9 [57.8, 74.8] | 59.7 [50.8, 67.9] | 61.3 [52.6, 69.3] |
| MLP | NUDZ | AUC | 80.2 [74.9, 84.9] | 75.4 [69.6, 81.0] | 76.1 [70.0, 81.5] | 74.7 [68.6, 80.4] | 66.6 [60.1, 72.9] | 74.2 [68.2, 80.0] |
| | | Sens. | 78.8 [73.4, 83.5] | 62.3 [55.1, 69.6] | 66.7 [59.5, 74.0] | 67.3 [60.4, 74.5] | 64.8 [57.6, 72.1] | 64.8 [57.6, 72.2] |
| | | Spec. | 70.2 [62.5, 77.5] | 72.1 [63.4, 80.7] | 71.2 [62.3, 79.5] | 71.2 [62.4, 79.6] | 60.6 [51.6, 70.2] | 72.1 [63.6, 80.5] |
| RowColCNN | IKEM | AUC | 87.2 [83.1, 90.9] | 78.1 [72.7, 83.3] | 78.2 [72.8, 83.2] | 77.9 [72.5, 83.1] | 69.4 [63.3, 75.3] | 79.5 [74.2, 84.4] |
| | | Sens. | 79.6 [73.9, 85.3] | 87.0 [81.2, 91.9] | 77.9 [71.1, 83.9] | 76.6 [69.9, 83.2] | 82.5 [76.4, 88.4] | 77.3 [70.4, 83.5] |
| | | Spec. | 78.7 [72.0, 84.7] | 50.8 [42.2, 59.2] | 63.7 [55.2, 71.7] | 64.5 [55.6, 72.8] | 39.5 [30.9, 48.2] | 64.5 [56.2, 72.7] |
| RowColCNN | NUDZ | AUC | 83.0 [78.0, 87.4] | 77.9 [72.1, 83.1] | 76.9 [71.1, 82.3] | 77.2 [71.4, 82.6] | 68.7 [62.0, 74.9] | 77.1 [71.1, 82.4] |
| | | Sens. | 82.8 [77.8, 87.4] | 71.0 [64.3, 77.6] | 70.4 [63.6, 77.5] | 56.8 [49.4, 64.3] | 77.2 [70.4, 83.2] | 69.1 [62.2, 76.0] |
| | | Spec. | 68.5 [60.5, 76.0] | 66.3 [57.0, 75.0] | 66.3 [57.3, 74.8] | 81.7 [73.9, 89.0] | 50.0 [40.7, 59.2] | 70.2 [61.3, 78.3] |
| *mfALFF* | | | | | | | | |
| RR | IKEM | AUC | 82.4 [77.6, 86.8] | 68.2 [61.8, 74.6] | 65.9 [59.2, 72.3] | 65.0 [58.2, 71.5] | 69.1 [62.8, 75.2] | n/a |
| | | Sens. | 75.5 [69.3, 81.2] | 77.9 [71.2, 84.5] | 69.5 [62.4, 76.7] | 70.1 [62.8, 77.4] | 71.4 [64.2, 78.5] | n/a |
| | | Spec. | 72.4 [64.9, 79.2] | 40.3 [31.7, 49.0] | 46.0 [37.4, 55.3] | 44.4 [35.8, 53.7] | 50.8 [42.5, 59.4] | n/a |
| RR | NUDZ | AUC | 78.0 [72.4, 83.1] | 66.0 [59.6, 72.6] | 68.8 [62.6, 75.1] | 68.3 [61.8, 74.6] | 64.5 [57.9, 70.9] | n/a |
| | | Sens. | 74.7 [68.4, 80.6] | 43.8 [36.5, 51.3] | 67.9 [60.5, 74.7] | 66.0 [58.6, 73.1] | 64.8 [57.8, 72.2] | n/a |
| | | Spec. | 62.7 [54.5, 71.1] | 77.9 [69.4, 85.7] | 56.7 [47.1, 66.3] | 57.7 [48.2, 67.3] | 55.8 [46.0, 65.6] | n/a |
| SVM | IKEM | AUC | 83.6 [79.0, 87.7] | 68.0 [61.7, 74.1] | 66.4 [59.9, 72.6] | 66.0 [59.6, 72.4] | 69.5 [63.2, 75.6] | n/a |
| | | Sens. | 76.4 [70.6, 82.0] | 80.5 [73.9, 86.5] | 73.4 [66.4, 80.2] | 74.0 [67.1, 80.6] | 78.6 [72.0, 84.9] | n/a |
| | | Spec. | 71.1 [63.7, 77.8] | 36.3 [28.0, 44.8] | 46.8 [38.3, 56.3] | 45.2 [36.6, 54.5] | 43.5 [34.5, 51.8] | n/a |
| SVM | NUDZ | AUC | 79.6 [74.3, 84.5] | 66.7 [60.1, 73.3] | 71.3 [65.2, 77.3] | 71.6 [65.5, 77.6] | 67.9 [61.4, 74.1] | n/a |
| | | Sens. | 76.0 [70.1, 81.5] | 48.1 [40.4, 55.7] | 69.1 [62.0, 75.8] | 69.1 [62.1, 75.8] | 69.8 [62.7, 76.5] | n/a |
| | | Spec. | 61.2 [52.9, 69.3] | 78.8 [70.8, 86.5] | 53.8 [44.2, 63.4] | 59.6 [50.0, 68.9] | 54.8 [45.0, 64.2] | n/a |
| MLP | IKEM | AUC | 83.7 [79.2, 87.9] | 61.5 [54.5, 68.0] | 58.3 [51.3, 65.3] | 60.2 [53.6, 66.7] | 53.8 [47.1, 60.6] | 52.3 [44.9, 59.4] |
| | | Sens. | 81.8 [76.4, 87.1] | 96.8 [93.4, 99.3] | 79.9 [73.0, 86.0] | 75.3 [68.2, 82.1] | 70.8 [63.4, 77.8] | 95.5 [91.9, 98.6] |
| | | Spec. | 72.6 [64.9, 79.4] | 6.5 [2.4, 11.1] | 31.5 [23.3, 39.5] | 33.9 [25.7, 42.6] | 40.3 [31.8, 49.1] | 16.1 [9.9, 22.8] |
| MLP | NUDZ | AUC | 76.3 [70.8, 80.9] | 60.8 [54.0, 67.5] | 70.9 [64.4, 77.0] | 69.4 [62.9, 75.6] | 61.2 [54.2, 67.9] | 68.6 [62.1, 74.6] |
| | | Sens. | 73.2 [67.3, 78.5] | 54.3 [46.2, 62.4] | 66.0 [58.5, 73.4] | 63.6 [55.9, 70.8] | 65.4 [58.2, 72.8] | 60.5 [53.0, 68.0] |
| | | Spec. | 67.3 [59.6, 75.0] | 56.7 [47.0, 66.7] | 64.4 [55.1, 73.9] | 64.4 [54.7, 74.0] | 48.1 [38.4, 57.6] | 66.3 [56.9, 75.7] |

Table 21: **Cross-site transfer on Schaefer100+Tian S2: AUC, sensitivity and specificity.** Area under the ROC curve (AUC), sensitivity (Sens.) and specificity (Spec.), in %, each the mean with its 95% bootstrap confidence interval in brackets, for cross-site transfer on functional connectivity (FC) and mfALFF features with the Schaefer100+Tian S2 atlas, by classifier and target site. Arms as in Table 16.

| Model | Target | Metric | Within-site | Direct | ComBat | CovBat | OT | ADDA |
|---|---|---|---|---|---|---|---|---|
| *Functional connectivity (FC)* | | | | | | | | |
| RR | IKEM | AUC | 91.6 [88.2, 94.5] | 86.4 [81.9, 90.6] | 86.3 [81.8, 90.5] | 87.0 [82.6, 91.0] | 81.6 [76.6, 86.3] | n/a |
| | | Sens. | 80.9 [75.3, 86.2] | 79.2 [72.7, 85.2] | 76.0 [69.0, 82.5] | 76.6 [69.7, 82.9] | 83.8 [77.7, 89.4] | n/a |
| | | Spec. | 84.0 [77.8, 89.7] | 77.4 [69.8, 84.1] | 82.3 [75.4, 88.5] | 82.3 [75.5, 88.2] | 63.7 [55.3, 72.3] | n/a |
| RR | NUDZ | AUC | 86.2 [81.7, 90.0] | 84.0 [78.9, 88.7] | 83.9 [78.9, 88.5] | 83.6 [78.5, 88.2] | 78.9 [73.3, 84.2] | n/a |
| | | Sens. | 76.7 [70.1, 82.2] | 73.5 [66.5, 80.1] | 75.9 [69.1, 82.4] | 74.7 [67.7, 81.3] | 74.7 [67.7, 81.2] | n/a |
| | | Spec. | 78.3 [70.5, 85.5] | 80.8 [73.4, 88.1] | 76.9 [68.7, 84.8] | 76.0 [67.6, 83.7] | 67.3 [58.3, 76.3] | n/a |
| SVM | IKEM | AUC | 91.6 [88.2, 94.5] | 84.8 [80.1, 89.1] | 84.9 [80.3, 89.3] | 86.3 [81.8, 90.3] | 76.7 [71.1, 82.0] | n/a |
| | | Sens. | 86.9 [82.1, 91.3] | 92.2 [87.9, 95.9] | 87.7 [82.1, 92.5] | 87.7 [82.4, 92.5] | 75.3 [67.9, 81.9] | n/a |
| | | Spec. | 80.3 [73.6, 86.4] | 58.1 [49.2, 66.7] | 71.0 [62.7, 78.8] | 71.8 [63.5, 79.7] | 62.1 [52.9, 70.1] | n/a |
| SVM | NUDZ | AUC | 85.6 [81.0, 89.6] | 83.5 [78.5, 88.2] | 83.6 [78.5, 88.3] | 83.7 [78.5, 88.4] | 79.2 [73.7, 84.5] | n/a |
| | | Sens. | 82.7 [76.9, 87.6] | 75.9 [68.9, 82.3] | 79.0 [72.6, 85.4] | 80.2 [74.1, 86.2] | 74.7 [68.2, 81.1] | n/a |
| | | Spec. | 71.3 [62.9, 79.4] | 74.0 [65.6, 82.1] | 70.2 [61.3, 78.8] | 71.2 [62.2, 79.6] | 65.4 [55.9, 74.3] | n/a |
| MLP | IKEM | AUC | 90.3 [86.9, 93.4] | 79.7 [74.3, 84.8] | 80.2 [74.7, 85.3] | 81.2 [76.0, 86.1] | 67.7 [61.4, 73.7] | 81.1 [75.8, 85.9] |
| | | Sens. | 83.9 [78.7, 88.8] | 89.0 [83.8, 93.7] | 81.2 [74.8, 86.6] | 81.2 [74.8, 86.7] | 68.2 [60.5, 75.2] | 80.5 [74.3, 86.2] |
| | | Spec. | 79.8 [73.2, 85.9] | 50.0 [41.0, 58.5] | 70.2 [61.9, 77.9] | 71.0 [62.8, 78.9] | 55.6 [46.8, 64.4] | 69.4 [61.5, 77.6] |
| MLP | NUDZ | AUC | 83.4 [78.5, 87.5] | 79.6 [74.0, 84.7] | 78.8 [72.9, 84.1] | 78.0 [72.1, 83.5] | 70.3 [63.8, 76.5] | 78.2 [72.2, 83.7] |
| | | Sens. | 79.6 [73.6, 84.9] | 68.5 [60.9, 75.2] | 72.8 [65.8, 79.4] | 69.8 [62.5, 76.4] | 61.7 [54.5, 68.5] | 72.2 [65.2, 78.8] |
| | | Spec. | 73.1 [65.2, 80.5] | 73.1 [64.0, 81.8] | 69.2 [60.2, 78.6] | 70.2 [61.1, 79.1] | 70.2 [61.4, 78.9] | 65.4 [56.0, 74.3] |
| RowColCNN | IKEM | AUC | 91.6 [88.5, 94.5] | 79.7 [74.4, 84.9] | 80.6 [75.3, 85.6] | 80.9 [75.7, 86.1] | 71.9 [65.6, 77.6] | 81.3 [76.1, 86.1] |
| | | Sens. | 83.1 [77.9, 88.1] | 84.4 [78.7, 90.0] | 74.0 [66.9, 80.5] | 72.1 [65.0, 78.9] | 84.4 [78.5, 89.9] | 72.7 [65.8, 79.6] |
| | | Spec. | 81.9 [75.7, 87.9] | 59.7 [50.8, 67.7] | 73.4 [65.2, 80.7] | 76.6 [69.2, 83.9] | 32.3 [24.2, 40.7] | 77.4 [69.8, 84.6] |
| RowColCNN | NUDZ | AUC | 84.6 [79.9, 88.6] | 80.0 [74.2, 85.0] | 81.3 [75.6, 86.1] | 79.2 [73.3, 84.4] | 72.0 [65.6, 78.0] | 80.6 [75.0, 85.5] |
| | | Sens. | 80.5 [74.8, 85.8] | 76.5 [69.9, 82.8] | 83.3 [77.5, 88.7] | 60.5 [52.8, 67.7] | 66.0 [58.4, 73.1] | 66.0 [58.6, 73.0] |
| | | Spec. | 74.2 [66.6, 81.1] | 67.3 [58.0, 76.4] | 64.4 [54.7, 73.3] | 80.8 [73.3, 88.3] | 69.2 [59.8, 77.7] | 79.8 [72.3, 87.2] |
| *mfALFF* | | | | | | | | |
| RR | IKEM | AUC | 80.4 [75.3, 85.1] | 68.6 [62.1, 74.6] | 71.8 [65.5, 77.4] | 72.4 [66.1, 77.9] | 68.6 [62.1, 74.7] | n/a |
| | | Sens. | 74.8 [68.4, 80.9] | 80.5 [74.0, 86.7] | 70.1 [62.6, 77.3] | 72.1 [64.6, 79.2] | 75.3 [68.2, 82.1] | n/a |
| | | Spec. | 70.0 [62.6, 77.1] | 50.8 [42.1, 59.6] | 54.8 [46.7, 63.5] | 53.2 [45.0, 61.9] | 52.4 [43.4, 60.9] | n/a |
| RR | NUDZ | AUC | 79.5 [73.8, 84.7] | 73.3 [67.1, 79.0] | 72.9 [66.9, 78.9] | 72.4 [66.4, 78.4] | 73.1 [66.9, 78.8] | n/a |
| | | Sens. | 76.2 [70.0, 82.0] | 29.6 [22.4, 36.6] | 72.8 [65.9, 79.5] | 73.5 [66.7, 80.1] | 67.3 [60.1, 73.9] | n/a |
| | | Spec. | 66.3 [58.4, 74.8] | 95.2 [90.7, 99.0] | 57.7 [48.2, 67.6] | 58.7 [49.0, 68.3] | 65.4 [55.7, 74.7] | n/a |
| SVM | IKEM | AUC | 81.4 [76.5, 85.9] | 69.6 [63.4, 75.6] | 73.1 [66.9, 78.8] | 73.7 [67.5, 79.3] | 70.1 [63.9, 76.2] | n/a |
| | | Sens. | 75.7 [69.4, 81.9] | 76.6 [69.5, 83.4] | 73.4 [66.0, 80.4] | 73.4 [65.6, 80.3] | 77.9 [71.3, 84.3] | n/a |
| | | Spec. | 71.6 [64.3, 78.3] | 53.2 [44.8, 62.5] | 54.0 [45.5, 62.9] | 55.6 [47.3, 64.5] | 50.0 [41.7, 59.1] | n/a |
| SVM | NUDZ | AUC | 79.0 [73.4, 84.1] | 74.0 [68.0, 79.5] | 74.8 [68.9, 80.4] | 74.5 [68.5, 80.3] | 73.2 [67.1, 79.0] | n/a |
| | | Sens. | 76.9 [71.3, 82.2] | 39.5 [32.1, 47.1] | 72.8 [66.1, 79.5] | 74.1 [67.5, 80.5] | 69.8 [62.3, 76.4] | n/a |
| | | Spec. | 63.1 [54.9, 71.1] | 85.6 [78.3, 92.1] | 59.6 [50.0, 69.2] | 59.6 [50.0, 69.2] | 61.5 [51.6, 71.0] | n/a |
| MLP | IKEM | AUC | 83.9 [79.3, 88.0] | 70.3 [63.7, 76.2] | 70.8 [64.3, 76.5] | 71.0 [64.6, 76.6] | 64.8 [58.1, 71.1] | 69.1 [62.7, 74.8] |
| | | Sens. | 80.6 [74.8, 86.0] | 97.4 [94.5, 99.4] | 89.0 [83.7, 93.5] | 89.6 [84.3, 94.0] | 67.5 [60.1, 74.8] | 94.2 [89.9, 97.5] |
| | | Spec. | 72.3 [65.3, 78.8] | 5.6 [1.9, 10.0] | 31.5 [23.6, 39.7] | 29.0 [21.2, 37.1] | 51.6 [42.9, 59.5] | 16.1 [9.9, 23.1] |
| MLP | NUDZ | AUC | 77.2 [71.9, 82.2] | 71.6 [65.6, 77.5] | 73.9 [67.8, 79.6] | 74.4 [68.3, 80.0] | 67.9 [61.4, 73.7] | 74.6 [68.7, 80.2] |
| | | Sens. | 74.7 [69.2, 79.8] | 38.3 [30.8, 45.9] | 71.6 [64.1, 78.3] | 74.1 [66.9, 80.7] | 73.5 [66.5, 80.0] | 56.2 [48.7, 63.9] |
| | | Spec. | 67.7 [60.5, 74.7] | 92.3 [86.8, 96.9] | 63.5 [53.9, 72.4] | 61.5 [51.5, 70.5] | 51.0 [41.7, 61.2] | 87.5 [80.6, 93.5] |

Table 22: **Cross-site transfer on Brainnetome246: AUC, sensitivity and specificity.** Area under the ROC curve (AUC), sensitivity (Sens.) and specificity (Spec.), in %, each the mean with its 95% bootstrap confidence interval in brackets, for cross-site transfer on functional connectivity (FC) and mfALFF features with the Brainnetome246 atlas, by classifier and target site. Arms as in Table 16.

| Model | Target | Metric | Within-site | Direct | ComBat | CovBat | OT | ADDA |
|---|---|---|---|---|---|---|---|---|
| *Functional connectivity (FC)* | | | | | | | | |
| RR | IKEM | AUC | 91.1 [87.7, 94.2] | 85.1 [80.4, 89.4] | 84.5 [79.9, 89.0] | 85.3 [80.7, 89.6] | 78.0 [72.6, 83.0] | n/a |
| | | Sens. | 81.6 [75.7, 87.0] | 83.8 [77.5, 89.1] | 76.6 [69.9, 83.1] | 77.3 [70.6, 83.7] | 77.9 [71.2, 84.3] | n/a |
| | | Spec. | 84.8 [78.8, 90.6] | 71.8 [64.3, 79.7] | 79.8 [72.8, 86.5] | 79.8 [72.8, 86.5] | 59.7 [50.8, 68.3] | n/a |
| RR | NUDZ | AUC | 84.9 [80.2, 88.9] | 81.7 [76.6, 86.5] | 81.4 [76.3, 86.3] | 81.5 [76.3, 86.4] | 75.1 [68.9, 80.7] | n/a |
| | | Sens. | 76.5 [70.4, 81.8] | 74.1 [67.3, 80.8] | 71.6 [64.7, 78.8] | 71.0 [64.2, 78.1] | 72.2 [64.7, 78.9] | n/a |
| | | Spec. | 76.0 [68.1, 83.4] | 72.1 [63.3, 80.6] | 75.0 [66.4, 83.3] | 75.0 [66.4, 83.5] | 61.5 [52.3, 70.7] | n/a |
| SVM | IKEM | AUC | 91.0 [87.6, 94.2] | 83.1 [78.4, 87.6] | 83.1 [78.4, 87.7] | 84.1 [79.4, 88.6] | 77.5 [71.9, 82.7] | n/a |
| | | Sens. | 87.9 [83.0, 92.2] | 93.5 [89.6, 97.2] | 85.7 [80.0, 91.0] | 87.7 [82.4, 92.5] | 81.2 [74.8, 87.1] | n/a |
| | | Spec. | 79.4 [72.5, 85.9] | 39.5 [31.2, 48.4] | 63.7 [55.6, 72.3] | 63.7 [55.1, 72.1] | 57.3 [48.7, 65.9] | n/a |
| SVM | NUDZ | AUC | 84.8 [80.1, 88.8] | 81.8 [76.7, 86.5] | 82.0 [76.8, 86.7] | 81.7 [76.5, 86.4] | 78.1 [72.0, 83.4] | n/a |
| | | Sens. | 86.8 [81.9, 91.2] | 75.3 [68.7, 81.8] | 79.6 [73.5, 85.7] | 79.0 [72.7, 85.3] | 77.2 [70.9, 83.3] | n/a |
| | | Spec. | 65.0 [56.8, 73.3] | 72.1 [63.2, 80.9] | 68.3 [59.4, 77.1] | 71.2 [62.3, 80.2] | 67.3 [58.0, 76.0] | n/a |
| MLP | IKEM | AUC | 89.8 [86.3, 93.0] | 78.6 [73.2, 83.7] | 79.9 [74.6, 84.9] | 80.4 [75.1, 85.4] | 64.3 [57.8, 70.7] | 79.5 [74.2, 84.7] |
| | | Sens. | 85.2 [80.0, 90.1] | 89.6 [84.8, 94.0] | 80.5 [73.7, 86.3] | 79.9 [73.3, 86.0] | 62.3 [55.3, 70.1] | 79.9 [73.5, 85.9] |
| | | Spec. | 80.6 [74.1, 86.8] | 40.3 [31.9, 49.3] | 69.4 [61.3, 77.1] | 65.3 [57.3, 73.3] | 65.3 [56.8, 73.3] | 68.5 [60.5, 76.2] |
| MLP | NUDZ | AUC | 81.3 [76.1, 85.8] | 76.8 [70.8, 82.3] | 77.7 [71.6, 83.1] | 76.6 [70.7, 82.3] | 67.5 [61.2, 73.5] | 75.2 [69.0, 81.2] |
| | | Sens. | 81.2 [75.7, 85.9] | 77.2 [70.5, 83.4] | 71.6 [64.6, 78.4] | 72.2 [65.4, 78.8] | 61.1 [53.7, 68.3] | 71.6 [64.6, 78.6] |
| | | Spec. | 70.6 [62.9, 78.0] | 59.6 [50.0, 68.9] | 68.3 [58.9, 77.7] | 72.1 [63.7, 81.0] | 65.4 [56.0, 74.3] | 67.3 [58.3, 76.7] |
| RowColCNN | IKEM | AUC | 90.9 [87.4, 94.0] | 78.7 [73.4, 84.0] | 77.8 [72.1, 83.2] | 77.4 [71.8, 82.9] | 67.1 [61.0, 73.2] | 78.6 [73.1, 83.9] |
| | | Sens. | 86.0 [81.4, 90.3] | 92.2 [87.6, 96.1] | 79.2 [72.5, 85.3] | 70.1 [62.8, 77.3] | 85.1 [79.4, 90.4] | 77.3 [70.4, 83.7] |
| | | Spec. | 81.8 [75.3, 88.0] | 41.9 [33.3, 50.8] | 67.7 [59.8, 75.8] | 73.4 [65.3, 81.4] | 25.8 [18.0, 33.6] | 70.2 [62.3, 78.0] |
| RowColCNN | NUDZ | AUC | 80.9 [75.8, 85.4] | 79.4 [73.6, 84.6] | 78.0 [72.2, 83.4] | 78.5 [72.6, 84.0] | 69.0 [62.2, 75.2] | 75.8 [69.6, 81.6] |
| | | Sens. | 84.0 [79.0, 88.2] | 78.4 [72.2, 84.7] | 72.8 [66.1, 79.5] | 56.2 [48.7, 63.7] | 78.4 [71.8, 84.5] | 64.2 [57.2, 71.4] |
| | | Spec. | 62.1 [54.4, 69.5] | 69.2 [60.5, 78.3] | 67.3 [58.8, 76.6] | 79.8 [72.4, 87.4] | 42.3 [32.6, 51.7] | 69.2 [60.7, 78.2] |
| *mfALFF* | | | | | | | | |
| RR | IKEM | AUC | 78.9 [73.6, 83.4] | 70.3 [63.9, 76.3] | 71.3 [64.9, 77.1] | 70.4 [64.0, 76.4] | 66.9 [60.5, 72.9] | n/a |
| | | Sens. | 72.7 [66.5, 78.5] | 94.2 [89.9, 97.5] | 74.7 [67.6, 81.3] | 74.0 [66.9, 80.7] | 71.4 [63.8, 78.2] | n/a |
| | | Spec. | 69.0 [61.7, 76.2] | 14.5 [8.4, 20.7] | 53.2 [44.7, 62.1] | 51.6 [43.0, 60.5] | 46.0 [36.7, 54.7] | n/a |
| RR | NUDZ | AUC | 79.5 [73.9, 84.5] | 64.7 [58.0, 71.0] | 66.2 [59.8, 72.6] | 67.9 [61.6, 74.2] | 62.6 [56.1, 69.0] | n/a |
| | | Sens. | 77.7 [72.0, 83.0] | 37.0 [30.1, 44.4] | 69.1 [61.9, 75.9] | 67.9 [60.7, 74.9] | 61.1 [53.7, 68.6] | n/a |
| | | Spec. | 66.9 [58.3, 75.1] | 81.7 [74.3, 88.9] | 51.9 [42.2, 61.3] | 56.7 [47.2, 66.3] | 56.7 [47.0, 65.7] | n/a |
| SVM | IKEM | AUC | 81.0 [76.1, 85.2] | 69.4 [63.1, 75.2] | 71.8 [65.6, 77.6] | 71.7 [65.3, 77.4] | 68.7 [62.4, 74.8] | n/a |
| | | Sens. | 72.5 [66.4, 78.0] | 96.1 [92.7, 98.8] | 75.3 [68.0, 81.9] | 76.0 [69.3, 82.5] | 74.0 [66.9, 80.8] | n/a |
| | | Spec. | 69.7 [62.6, 76.5] | 8.1 [3.5, 13.2] | 54.0 [45.3, 63.0] | 51.6 [42.3, 60.7] | 48.4 [39.7, 57.0] | n/a |
| SVM | NUDZ | AUC | 80.0 [74.6, 84.7] | 62.1 [55.0, 68.7] | 68.2 [61.6, 74.5] | 70.4 [63.9, 76.5] | 61.3 [54.5, 68.1] | n/a |
| | | Sens. | 77.8 [72.3, 83.2] | 40.1 [32.7, 47.7] | 66.7 [59.1, 73.9] | 64.2 [57.1, 71.5] | 66.7 [59.6, 73.8] | n/a |
| | | Spec. | 64.2 [56.2, 72.0] | 77.9 [69.8, 85.6] | 55.8 [46.0, 64.9] | 55.8 [46.5, 65.2] | 48.1 [38.3, 58.0] | n/a |
| MLP | IKEM | AUC | 82.2 [77.4, 86.3] | 59.8 [52.9, 66.2] | 61.4 [54.7, 67.6] | 61.1 [54.1, 67.4] | 60.0 [53.4, 66.6] | 54.6 [47.8, 61.3] |
| | | Sens. | 79.2 [73.4, 84.7] | 96.1 [92.7, 98.8] | 85.1 [79.2, 90.3] | 83.8 [77.8, 89.2] | 76.6 [69.6, 82.8] | 89.0 [83.7, 93.6] |
| | | Spec. | 69.0 [62.1, 75.7] | 5.6 [1.7, 9.8] | 25.8 [18.4, 33.6] | 28.2 [20.5, 36.4] | 30.6 [22.6, 38.6] | 14.5 [8.8, 20.8] |
| MLP | NUDZ | AUC | 76.8 [71.4, 81.9] | 63.7 [56.9, 70.0] | 69.7 [63.4, 75.6] | 70.7 [64.3, 76.5] | 57.8 [50.6, 64.5] | 69.3 [62.9, 75.0] |
| | | Sens. | 78.3 [72.9, 83.4] | 48.1 [40.1, 56.0] | 66.7 [59.5, 73.7] | 67.9 [60.5, 75.0] | 71.6 [64.4, 78.4] | 58.6 [51.3, 66.3] |
| | | Spec. | 62.5 [55.0, 69.8] | 76.9 [68.9, 84.6] | 58.7 [48.9, 67.7] | 58.7 [49.1, 67.8] | 44.2 [34.9, 53.5] | 72.1 [63.2, 80.2] |
